# Androgen Receptor SUMO site mutations ameliorate toxicity in *Drosophila* models of SBMA

**DOI:** 10.64898/2026.10.23.761456

**Authors:** Kristin Richardson, Matthew V. Prifti, Truman Kula, Fatimah H. Hussein, Sokol V. Todi

## Abstract

Spinal and bulbar muscular atrophy (SBMA) is an X-linked neurodegenerative disorder caused by a CAG trinucleotide repeat expansion in the androgen receptor (AR) gene that produces a toxic, polyglutamine-expanded protein. SBMA is part of a broader class of polyglutamine (polyQ) diseases characterized by protein misfolding, aggregation, and progressive neuronal dysfunction, for which no disease-modifying therapies exist. Post-translational modification of AR, including SUMOylation, is an important regulator of its transcriptional activity, stability, and cellular localization. Disease protein modification by SUMO is implicated in various neurodegenerative diseases and can influence protein aggregation and toxicity in polyQ disorders. Prior studies suggest that preventing AR SUMOylation positively modifies disease phenotypes. Here, we generated *Drosophila* models expressing polyQ-expanded human AR in which the canonical SUMO acceptor lysine sites were mutated into arginines to prevent SUMOylation of AR. Using tissue-specific expression in neurons and muscle, we assessed effects on lifespan, motor behavior, AR localization, and gene expression. Mutation of AR SUMOylation sites leads to improvements in longevity when expressed in muscle. Effects on motor performance were more limited. Transcriptomic analysis revealed tissue-specific effects of AR SUMO-site mutation, with enrichment of immune, metabolic, and defense-related processes in heads, but limited enrichment in thoraces. In both tissues, a subset of genes altered by AR SUMO-site mutation showed expression changes opposing those associated with polyQ expansion. These findings support a modulatory role for SUMOylation in SBMA pathogenesis and highlight post-translational regulation of AR as a potential therapeutic avenue requiring further investigation.

## INTRODUCTION

Spinal and bulbar muscular atrophy (SBMA, also known as Kennedy’s disease), is an X-linked, recessive neurodegenerative disorder caused by a CAG trinucleotide repeat expansion in the polyglutamine (polyQ) tract of the androgen receptor (AR) gene^1^. SBMA belongs to a family of polyQ disorders, each caused by expansion of a polyQ tract within disease-specific proteins. Other members of this family include Huntington’s disease, dentatorubral–pallidoluysian atrophy, and several forms of spinocerebellar ataxia (types 1, 2, 3, 6, 7, and 17)^2,3^. Although SBMA is not inherently fatal, it leads to severe neuromuscular deterioration that impairs strength, mobility, and bulbar functions such as swallowing and speech, resulting in a marked decline in quality of life^4^. Despite its clinical impact, there are currently no approved therapies that can prevent disease progression.

AR is a ligand-dependent transcription factor whose activity is regulated via multiple post-translational events^5^. SUMOylation is one such conserved post-translational modification that acts as an important regulator of AR function^5,6^. It involves the covalent attachment of small ubiquitin-like modifier (SUMO) proteins to specific lysine residues on target proteins, thereby altering their activity within the cell^7,8^. SUMOylation participates in a wide range of intra- and extra-nuclear biological processes, including transcription, nuclear transport, synaptic transmission, neuronal excitability and cellular responses to stress^9,10^. It also plays an important role in cellular protein quality control^11^. Similar to ubiquitination, SUMOylation requires sequential enzymatic events for its activation and conjugation onto substrate proteins^12,13^. Once activated, SUMO proteins can be added to substrates either as single monomers or as polymeric chains and can be removed by de-SUMOylating proteases^7,12–14^. The (dys)regulation of SUMOylation has been implicated in a variety of neurodegenerative diseases^15^, including Parkinson’s disease^16,17^, Alzheimer’s disease^18,19^, and several polyQ disorders^20–23^, highlighting its relevance to protein aggregation-associated neurodegeneration.

SUMOylation has a well-established inhibitory effect on AR transcriptional activity. In the context of SBMA, preventing SUMOylation of disease-causing, polyQ-expanded AR, and therefore removing the inhibitory effect of SUMO, enhances transcriptional activity in ARQ112 SBMA cell models and partially restores expression patterns in AR-responsive genes^22^. Similar transcriptional effects, in addition to significant improvement in lifespan, were also observed in SBMA mice when AR could not be SUMOylated, despite comparable levels of polyQ-expanded AR expression and aggregation^22^. In contrast to the beneficial effects of targeted disruption, broader disruption of SUMO activation pathways through the expression of a non-functional Uba2 SUMO-activating enzyme, together with a truncated form of polyQ-expanded AR, exacerbated neurodegeneration in the *Drosophila* eye^24^. Conversely, broadly enhancing SUMOylation via SUMO protein overexpression reduced AR aggregation in cell models of SBMA in a manner that was dependent on SUMOylation of AR itself and not of other SUMO substrates^25^. Collectively, these studies indicate that SUMO plays a complex modulatory role in SBMA, with effects that are likely context- and function-dependent, underscoring the need for further investigation into how SUMOylation of polyQ AR influences pathogenesis.

While five SUMO proteins exist in humans^7,26^, *Drosophila* have one, highly conserved, SUMO ortholog, Smt3^27^, which serves as the sole mediator of SUMOylation. Smt3 is most functionally comparable to the SUMO 2/3 subgroup in humans^28^, with functional relevance in diverse biological processes including neural development^29^, inflammatory signaling^30,31^, and transcriptional activity^32–34^. However, unlike SUMO 2/3 in vertebrates, Smt3 is unable to form poly-SUMO chains, and loss of Smt3 cannot be rescued by human SUMO expression, highlighting some evolutionary divergence despite high sequence homology^35^. Still, the conservation of SUMO functions between flies and mammals, combined with the genetic tractability of *Drosophila*, provides a strong experimental framework for investigating the role of SUMOylation in SBMA pathology.

To contribute to the growing literature aimed at better understanding these mechanisms, we generated and characterized a new SUMO mutant *Drosophila* model of SBMA, providing an additional in vivo system to dissect how SUMOylation impacts AR function and associated phenotypes. We modified the human AR sequences used in our previously published SBMA models^36^ to mutate SUMO acceptor site lysine residues to arginine, consequently disrupting SUMOylation, and assessed the effects of these changes on SBMA pathology. We examined changes in longevity, motility, AR localization and gene expression in neurons and in muscles. We observe that SUMO-site mutation of polyQ AR extends lifespan when expressed in muscle, without obvious alterations to AR localization or aggregation, and produces tissue-dependent transcriptional changes that largely oppose the transcriptional consequences of polyQ expansion. A further understanding of how SUMOylation modulates AR function in SBMA may reveal novel therapeutic targets to lessen pathogenic changes in disease.

## RESULTS

### Generation of polyQ-expanded AR-SUMO-site-mutated Drosophila lines

Our previous work established a robust model of SBMA in *Drosophila* with strong disease phenotypes independent of ligand binding^36^. To investigate the contribution of SUMOylation to SBMA pathology, we modified the full-length human AR cDNA sequence used to generate these established lines by substituting the SUMO acceptor lysine residues with arginine residues (Figure 1A and supplemental figure 1A). Single nucleotide substitutions from A to G (A**A**G to A**G**G) were introduced to generate the SUMO-null construct (supplemental figure 1B). Human AR has two established SUMO acceptor sites located at K388 and K521 in the reference sequence used to generate our models^37,38^. In our ARQ112 modified sequence, these sites are shifted to K478 and K611 due to the expanded polyQ region. Stable transgenic fly lines were generated using PhiC31-dependent integration into a “safe-harbor” locus on chromosome 3. Integration of the correct sequence was verified by Sanger sequencing, and expression of AR transcript and protein was confirmed by Western blot and qPCR, respectively (Figure 1B, and 1C).

**Figure 1:**
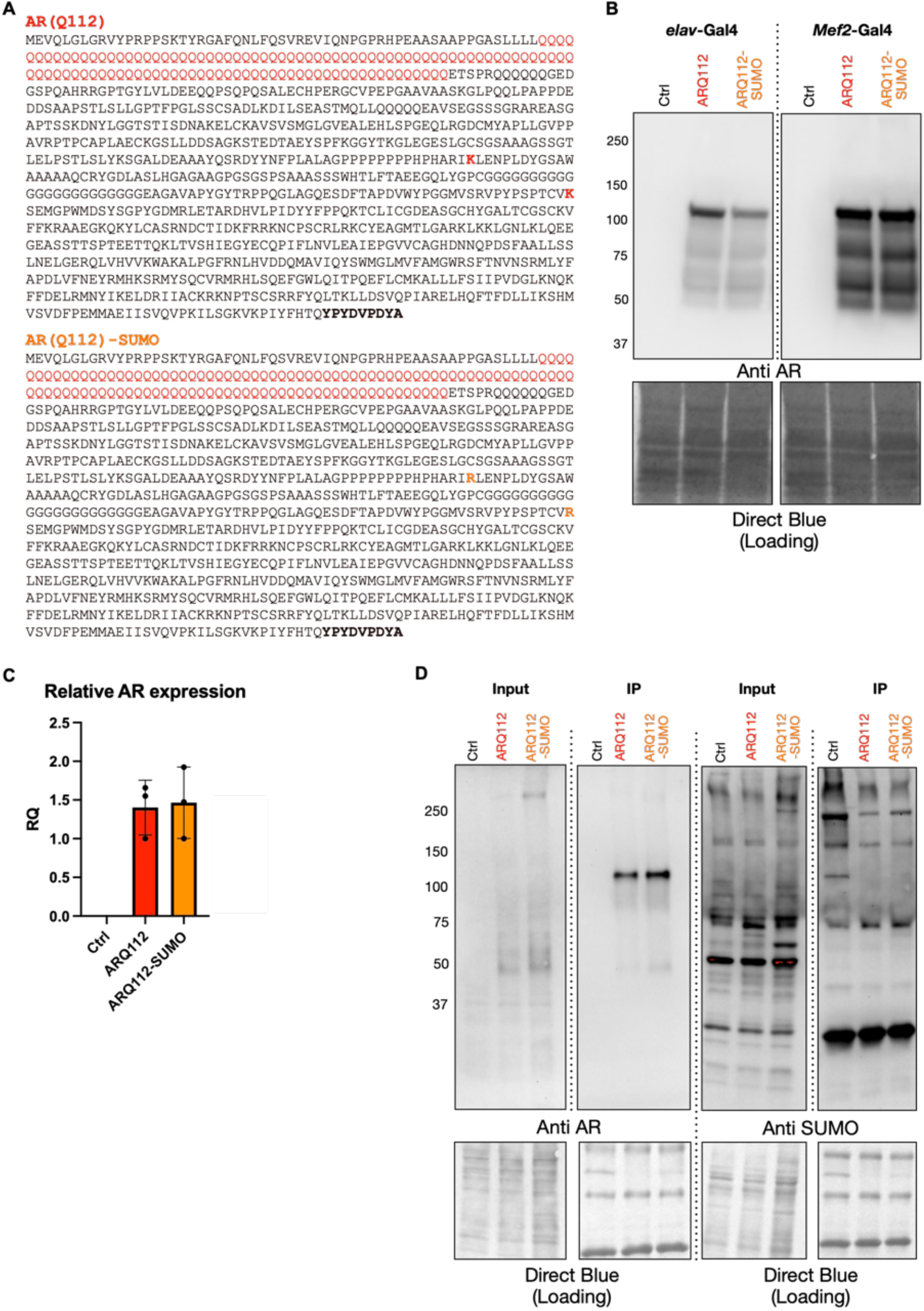
Generation of SUMO-site mutant SBMA flies. A) Amino acid sequences of the human AR proteins used. ARQ112: unmodified AR protein, with lysine (K) residues at established SUMOylation sites highlighted in red. ARQ112-SUMO: SUMO-site mutated AR protein, with substituted arginine (R) residues highlighted in orange. Bold: HA epitope tag. B) Western blot from adult female flies expressing the noted transgenes. *elav*-Gal4: pan-neuronal expression. *Mef2*-Gal4: muscle expression. C) qRT-PCR of AR transcript levels in adult flies expressing ARQ112 or ARQ112-SUMO in muscle. No amplification was detected in the driver-only control (expression below the detection threshold); the value is displayed as zero for visualization purposes only. N = 3 biological replicates per group. Statistics: Welch’s unpaired t-test, mean ± SD; not significant (p = 0.90). D) Western blots from anti-HA immunoprecipitation of HA-tagged ARQ112 and ARQ112-SUMO proteins expressed in adult female flies. Blots were run on 4-20% gels and probed with anti-AR and anti-SUMO antibodies. N=1 independent lysate per condition. IP: immunoprecipitated fraction. Input: lysate prior to immunoprecipitation.

To determine whether human AR is SUMOylated by endogenous *Drosophila* SUMO proteins, and to assess whether the introduced SUMO mutations prevented SUMO conjugation to AR, we performed stringent immunoprecipitations using the c-terminal HA tag to isolate AR (Figure 1D). While successful pull-down of AR was confirmed in IP samples, we were unable to biochemically detect SUMO-conjugated species. This is likely due to both the limited number of available reagents targeting SUMO in the fly, including a dearth of antibodies, and the transient nature of protein SUMOylation.

### Mutating SUMOylation sites of PolyQ-expanded AR in muscle extends lifespan in Drosophila

Previous studies in mammalian cell culture and in mice have shown reduced toxicity across multiple disease parameters following mutation of SUMO acceptor sites on polyQ AR^22,25^. To test this in our fly models, we examined longevity and motility outcomes in ARQ112-SUMO flies. Since our SBMA model is not hormone-dependent and was originally characterized in female flies^36^, we conducted all experiments in females.

Because toxicity in SBMA has been attributed to pathology in both neurons and muscle^39,40^, we utilized the GAL4/UAS system^41^ to drive expression of our constructs in these tissues. While there was no difference in longevity between ARQ112 and ARQ112-SUMO when expressed in neurons (Figure 2A, top), expression of ARQ112-SUMO in muscle significantly improved longevity (Figure 2B, top). Of note, male ARQ112-SUMO flies also gained improvements to longevity when expressed in muscle (Supplemental Figure 2). We did not observe any significant differences in motility with expression in either tissue, whether examining rate of decline in longitudinal performance (Figure 2A and B, bottom) or week-to-week changes in overall performance (Supplemental Figure 3). Levels of ARQ112 protein were also unchanged in both tissues (Figure 2C; neurons, and 2D; muscle), suggesting that the longevity improvement observed in muscle was not due to marked changes in protein levels.

**Figure 2:**
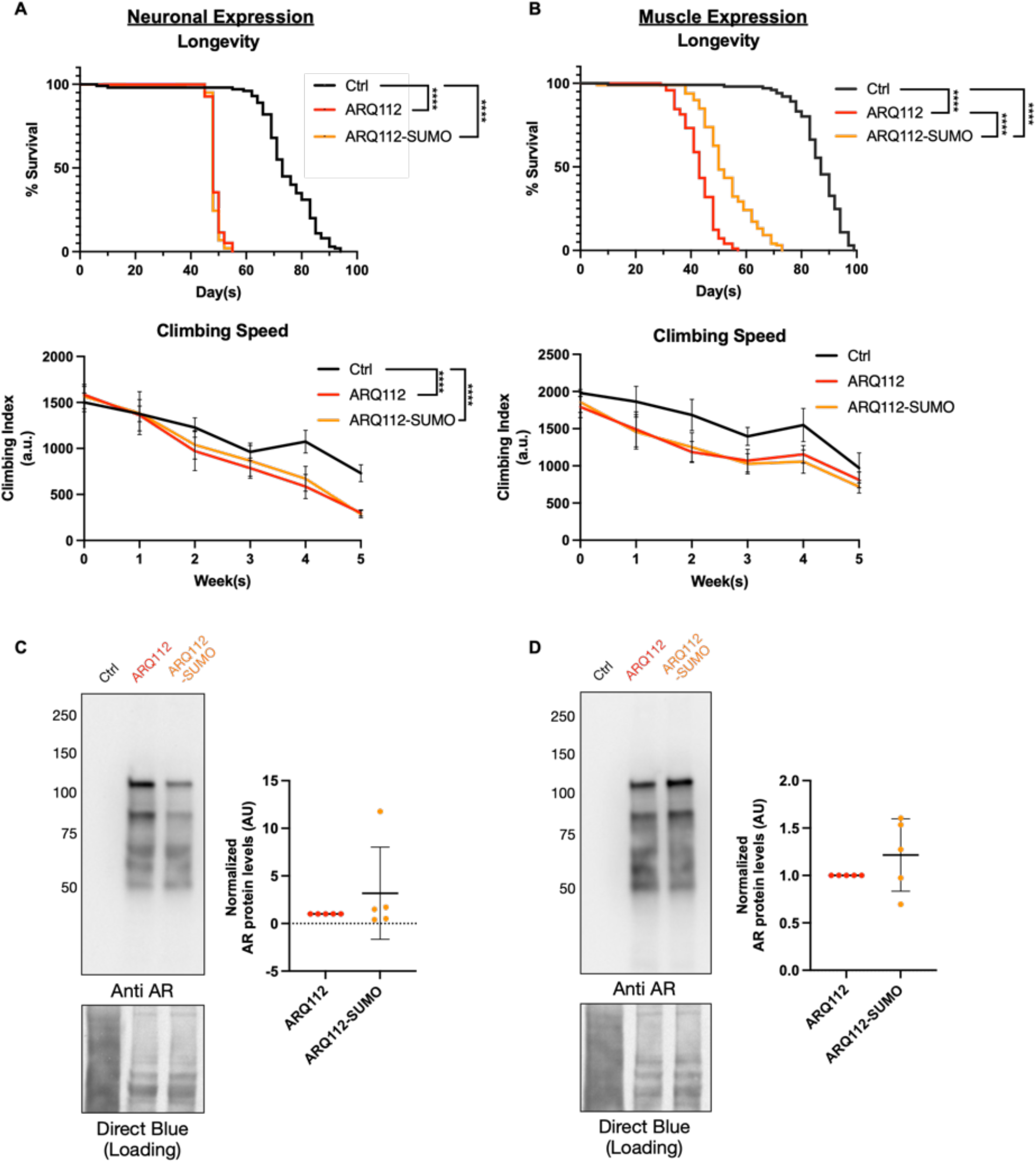
Effects of ARQ112-SUMO expression on longevity, motility and AR protein levels. A) Longevity (top) and motility (bottom) outcomes with neuronal expression of noted transgenes. B) Longevity (top) and motility (bottom) outcomes with muscle expression of noted transgenes. For both (A) and (B), N=100 flies per group. Statistics: Log-rank tests, ****: p<0.0001 (longevity); linear regression comparing slopes, mean ± SD, ****: p<0.0001 (motility). C) Western blot from 3-day old flies expressing the noted AR proteins in neurons. D) Western blot from 3-day old flies expressing the noted AR proteins in muscle. For both (C) and (D), N=5 independent lysates per condition were prepared, electrophoresed and quantified. Blots were run on 4-20% gels and ARQ112-SUMO protein levels were normalized to those of matched ARQ112 samples. Statistics: not significant by Wilcoxon signed-rank test, mean ± SD.

Since rapid iterative negative geotaxis (RING)-based motility assays strictly measure how quickly flies initiate climbing behaviors following a negative geotaxis stimulus within a short window of time, we wondered whether mobility phenotypes might be missed using this assay alone. We therefore also measured longer-duration timed mobility. We again observed no lasting significant differences in mobility between genotypes. However, ARQ112-SUMO flies performed statistically better than ARQ112 at day 7 in neurons, and statistically worse at day 35 in neurons (Figure 3A) and day 21 in muscle (Figure 3B). In each case, this difference was no longer observed by the following week. Supplemental video 1 provides a visual example of climbing ability across genotypes at day 35.

**Figure 3:**
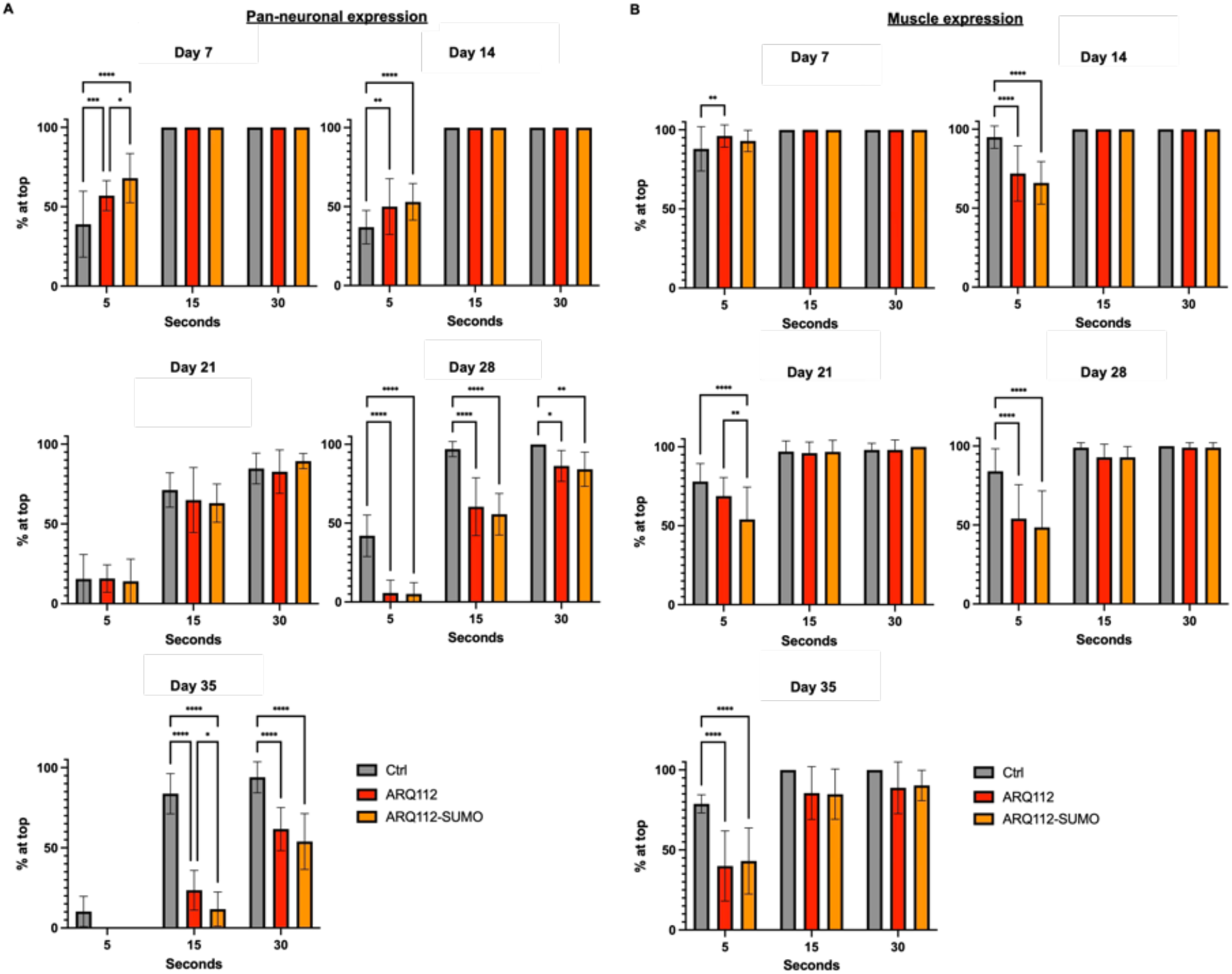
Effects of ARQ112-SUMO expression on time-dependent climbing performance. Climbing ability at 5, 15 and 30 seconds in adult flies expressing the noted transgenes in (A) neurons or (B) muscle. N=200 flies per group. Statistics: 2-way ANOVA + Tukey post-hoc, mean ± SD. *: p<0.05, **: p<0.01, ***: p<0.001, ****: p<0.0001.

Overall, these data support a role for AR SUMOylation in modulating SBMA toxicity in a tissue-dependent manner, since disruption of AR SUMOylation sites improved longevity when expressed in muscles but not in neurons. This improvement occurred without substantial changes to AR protein levels, suggesting that the improvement is likely not a result of reduced or improved clearance of toxic AR protein. Instead, SUMOylation status may influence downstream mechanisms that contribute to disease progression.

### SUMO-site mutation does not alter AR localization or aggregation in neurons or muscle

Since improved longevity in ARQ112-SUMO flies was not explained by differences in AR protein abundance, we next examined whether disruption of AR SUMOylation altered other disease-associated properties of polyQ-AR, including nuclear localization and aggregation. We first examined dissected adult brain tissue from young and aged flies to assess protein localization patterns following pan-neuronal expression of our ARQ112 constructs (Figure 4).

**Figure 4:**
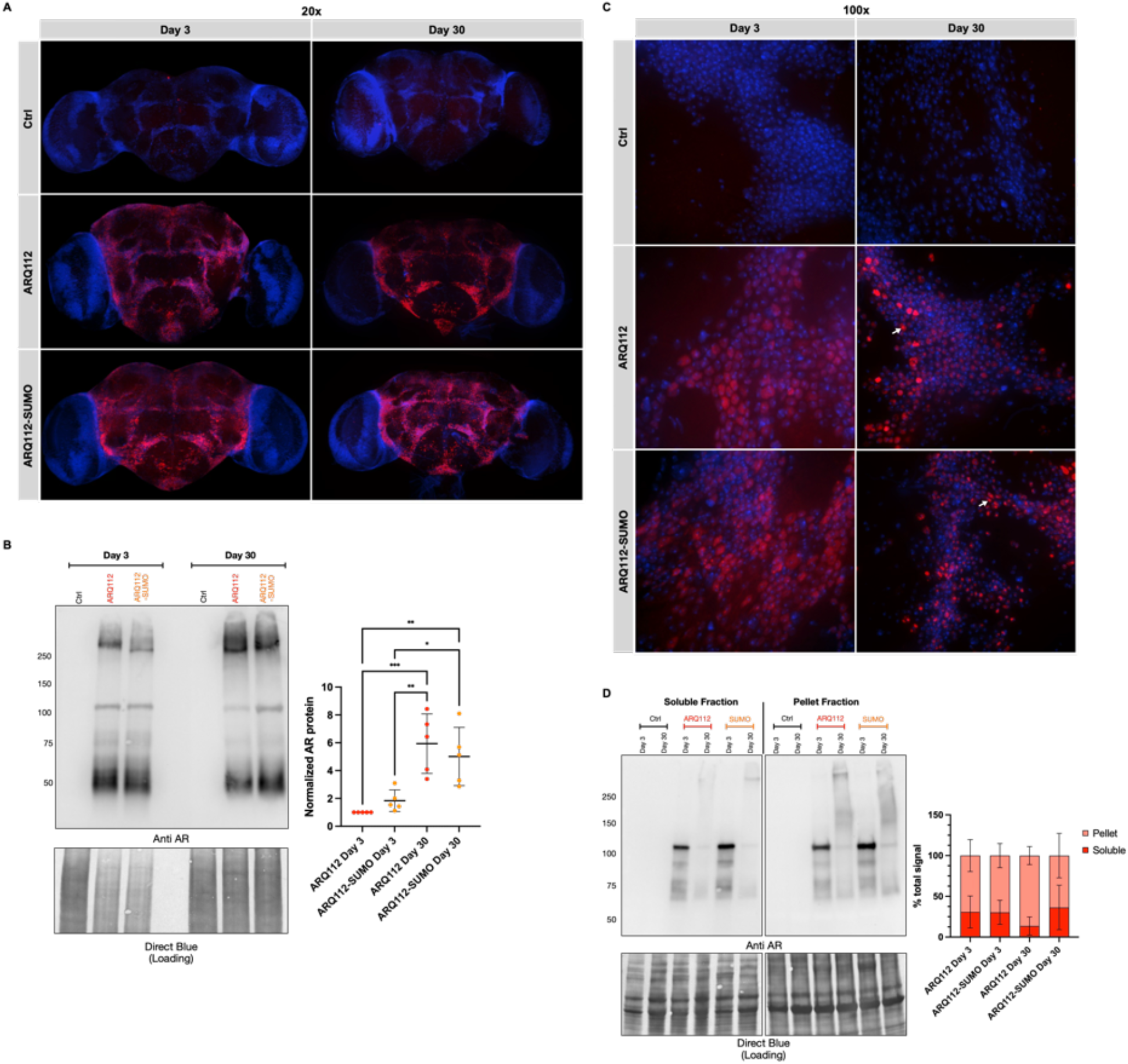
Effects of ARQ112-SUMO expression in neurons on AR protein localization, accumulation and aggregation. A) Representative immunofluorescence images of dissected adult fly brains at day 3 and day 30. 20x magnification, Red: anti-AR, Blue: DAPI. B) Western blot from adult flies expressing the noted AR transgenes at day 3 and day 30. N=5 independent lysates per condition. Blots were run on 7.5% gels and AR protein levels were normalized to matched day 3 ARQ112 samples. Statistics: 2-way ANOVA + Tukey post hoc, mean ± SD. *: p<0.05, **: p<0.01, ***: p<0.001. C) Representative immunofluorescence images of dissected adult fly brains at day 3 and day 30. 100x magnification, Red: anti-AR, Blue: DAPI, white arrows: cells where AR is predominantly nuclear, with AR staining obscuring DAPI signal. D) Western blots of soluble and pellet fractions from adult flies at day 3 and day 30. N=5 independent lysates per condition. Blots were run on 4-20% gels and quantified as a percentage of the total AR signal. Statistics: not significant by unpaired Welch’s t-test, mean ± SD.

In whole-brain tissue, the overall distribution of AR signal did not visually differ between ARQ112 or ARQ112-SUMO conditions at either day 3 or day 30 (Figure 4A). However, Western blot analysis of whole fly lysates revealed an age-dependent increase in AR protein levels in both conditions, especially in the higher portions of the blot (Figure 4B), suggesting increased aggregation with age. To more closely examine potential changes in subcellular localization of AR, we turned to higher-magnification images to assess nuclear versus cytoplasmic distribution in individual cells (Figure 4C). At day 3, AR signal appears to be predominately cytoplasmic. By day 30, AR signal displayed a more punctate staining pattern, which in some cells appeared to obscure the DAPI nuclear signal altogether (Figure 4B, white arrows). These patterns were observed irrespective of SUMO-site mutation status. Subcellular fractionation and subsequent western blot showed no significant differences at either day 3 or day 30 in the cytoplasmic or nuclear distribution of ARQ112 versus ARQ112-SUMO (supplemental figure 4A). However, AR protein was difficult to detect in day 30 fractions, particularly in the nuclear fractions, limiting our ability to confidently assess AR localization using this method.

To examine AR aggregation with pan-neuronal expression, we conducted western blot analysis of soluble and insoluble (pellet) fractions (Figure 4D). In soluble fractions, both ARQ112 and ARQ112-SUMO displayed a distinct AR band at day 3, whereas day 30 samples shifted to a smear of higher molecular weight species, consistent with increased aggregation-prone AR. However, quantification of total soluble AR revealed no significant difference between day 3 and day 30 samples in either ARQ112 or ARQ112-SUMO conditions. Insoluble fractions followed the same patterns, with no statistical differences observed. A trend toward a larger percent of AR protein in the insoluble fraction was observed at day 30 in ARQ112 flies, however this did not reach statistical significance (p=0.33).

Given the muscle-specific improvement in longevity observed in ARQ112-SUMO flies, we next assessed AR localization and aggregation following muscle-specific expression of our ARQ112 constructs (Figure 5). In both ARQ112 and ARQ112-SUMO tissues, we observed no visual differences in AR signal localization between ARQ112 and ARQ112-SUMO at either day 3 or day 30 (Figure 5A), consistent with subcellular fractionation blots showing no significant differences in cytoplasmic versus nuclear AR protein distribution (supplemental figure 4B). However, a striking age-dependent change in AR staining pattern was observed in both conditions. At day 3, AR appeared as numerous small puncta, while at day 30 the signal was concentrated in fewer, much larger punctate structures (Figure 5A; 100x zoom). Western blot analysis using whole fly lysates again showed no significant increase in AR protein levels between day 3 and day 30 samples under either condition (Figure 5B). Analysis of soluble and insoluble fractions revealed a trend toward increased higher molecular weight species of AR at day 30 in both ARQ112 and ARQ112-SUMO samples, however none of these changes reached statistical significance (p=0.24 and p=0.08, respectively) (Figure 5C; All biological replicates of the soluble and pellet fractionation experiments are shown in supplemental figure 5).

**Figure 5:**
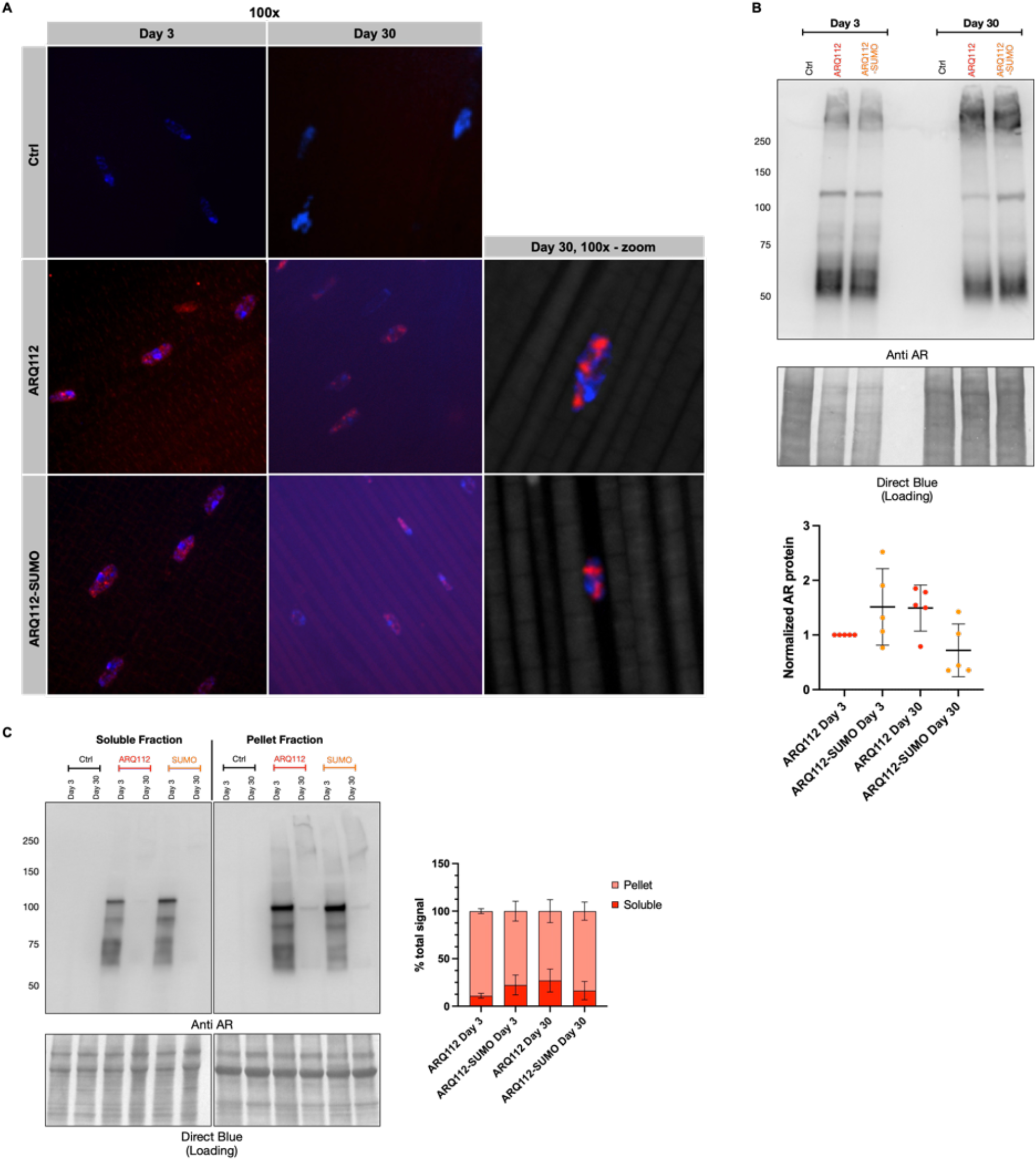
Effects of ARQ112-SUMO expression in muscle on AR protein localization, accumulation and aggregation. A) Representative immunofluorescence images of dissected dorsal longitudinal flight muscle at day 3 and day 30. 100x magnification, Red: anti-AR, Blue: DAPI. B) Western blot from adult flies expressing the noted AR transgenes at day 3 and day 30. N=5 independent lysates per condition. Blots were run on 7.5% gels and AR protein levels were normalized to matched day 3 ARQ112 samples. Statistics: not significant by 2-way ANOVA + Tukey post hoc, mean ± SD. C) Western blots of soluble and pellet fractions from adult flies at day 3 and day 30. N=5 independent lysates per condition. Blots were run on 4-20% gels and quantified as a percentage of the total AR signal. Statistics: not significant by unpaired Welch’s t-test, mean ± SD.

Together, these data suggest that AR protein localization and aggregation are not markedly altered in ARQ112-SUMO conditions in neurons or in muscle compared to ARQ112 alone. Interestingly, and consistent with our previously published results, most of the polyQ-AR signal in both tissues seems to be concentrated at the nucleus, despite the lack of ligand. The same was observed for wild-type AR (ARQ20); however, the signal in this case appears more perinuclear than intranuclear (Supplemental Figure 6). Further studies will be needed to determine the mechanistic basis for ligand-independent nuclear enrichment of AR in our model.

### Gene expression analysis reveals tissue-specific transcriptional changes in ARQ112-SUMO flies

In the absence of apparent changes to AR protein expression, aggregation, or localization in ARQ112-SUMO flies, we sought to determine whether disrupting AR SUMOylation led to downstream changes in gene expression that could contribute to muscle-specific improvements in longevity.

To address this, we conducted RNA-sequencing of female flies expressing no AR transgene (Ctrl; driver outcrossed to the background control line, w^1118^), wild-type AR (ARQ20), ARQ112 or ARQ112-SUMO in either neurons (heads) or muscle (thoraces) to examine tissue-specific transcriptional changes. Flies were aged 7 days to examine gene expression prior to disease onset, allowing us to identify transcriptional changes associated with expression of our transgenes rather than as consequences of later-stage disease progression. Expression of each transgene was confirmed by western blot (Supplemental Figure 7). Principal component analysis (PCA) was used to assess the global transcriptional variation among samples (Supplemental Figure 8). In both heads and thoraces, Ctrl samples separated from AR-expressing genotypes, whereas ARQ20, ARQ112 and ARQ112-SUMO samples showed more substantial overlap. One ARQ112 thoracic sample (ARQ112 sample 3) was identified as an outlier based on a formal statistical assessment and was therefore excluded prior to all subsequent analyses.

We next examined differential gene expression to identify transcriptional changes associated with exogenous expression of wild-type AR, polyQ-expansion of AR, and disruption of AR SUMOylation sites. Of particular interest to us was whether ARQ112-SUMO expression altered transcriptional changes associated with ARQ112 expression and whether these effects differed between neuron-and muscle-enriched tissues. Significant differentially expressed genes (DEGs) identified across each comparison are summarized in Venn diagrams (Figure 6A), with individual genes and associated expression changes for each comparison provided in supplemental tables 1 and 2. Expression of ARQ20 resulted in the greatest number of DEGs relative to Ctrl, with 346 identified in heads and 1,099 in thoraces. In comparison, ARQ112 resulted in 71 DEGs in heads and 70 in thoraces, while ARQ112-SUMO resulted in 61 DEGs in heads and 38 in thoraces.

A subset of genes altered by ARQ112-SUMO overlapped with those altered by ARQ112. In heads, 29 of the 61 ARQ112-SUMO associated DEGs were also differentially expressed between ARQ112 and ARQ20, while 18 were exclusive to the ARQ112-SUMO versus ARQ112 comparison. Similarly, 15 of the 38 ARQ112-SUMO-associated DEGs overlapped with the ARQ112 versus ARQ20 comparison in muscle, while 12 were exclusive to the ARQ112-SUMO versus ARQ112 comparison.

The distribution and intersections of these DEGs across all six tissue and genotype comparisons are shown by UpSet analysis in supplemental figure 9. Between heads and thoraces, unsurprisingly, the largest cross-tissue overlap existed between the ARQ20 versus Ctrl comparisons, which exclusively shared 93 DEGs. Comparatively, only two DEGs, CG12716 and the pseudogene CR40190, were shared exclusively between the ARQ112 versus ARQ20 comparisons, while only one gene, CG30184, was shared exclusively between the ARQ112-SUMO versus ARQ112 comparisons. Three genes were represented across all six comparisons. Together, these results suggest that the transcriptional changes associated with ARQ112 and ARQ112-SUMO expression are predominantly tissue-specific.

To determine whether these transcriptional changes were associated with specific biological processes, we next performed gene ontology (GO) and KEGG pathway enrichment analysis (Figure 6B and C). In heads, enriched GO terms were predominantly associated with immune and defense responses to bacteria and fungi, with several of these terms identified in both ARQ112 and ARQ112-SUMO samples but not in ARQ20 versus Ctrl. In contrast, ARQ20 expression in thoraces resulted in a strong enrichment for processes related to nuclear division, chromosomal segregation, mitotic cell cycle regulation and respiratory electron transport. Little to no GO enrichment was observed for ARQ112 or ARQ112-SUMO samples in thoraces. KEGG pathway analysis showed similar tissue- and genotype-dependent differences. ARQ20 expression in thoraces was strongly associated with enrichment of oxidative phosphorylation, as well as pathways related to DNA replication and repair. Fewer pathways were identified across the remaining comparisons. ARQ112-SUMO expression in heads was associated with enrichment of metabolic pathways, while ARQ112 expression in thoraces was associated with Toll and Imd signaling and neuroactive ligand-receptor interaction. No significant KEGG pathways were identified for ARQ112-SUMO relative to ARQ112 in thoraces.

To further examine gene expression patterns across these comparisons, we generated volcano plots to visualize the magnitude and direction of expression changes among the differentially expressed genes (Figure 7). The top 10 genes based on statistical significance are labeled for each comparison. Expression of ARQ20 relative to Ctrl in both tissues produced the broadest transcriptional response, with differentially expressed genes distributed in both directions. While many of the most statistically significant changes were observed in heads, thoraces showed larger fold changes in gene expression despite generally lower statistical significance. In thoraces, target of brain insulin (tobi) stood out as both the most statistically significant DEG and the gene with the largest fold change in the ARQ20 versus Ctrl comparison. ARQ112 versus ARQ20 and ARQ112-SUMO versus ARQ112 showed narrower distributions overall, although individual genes still showed large changes in expression in both directions.

**Figure 6:**
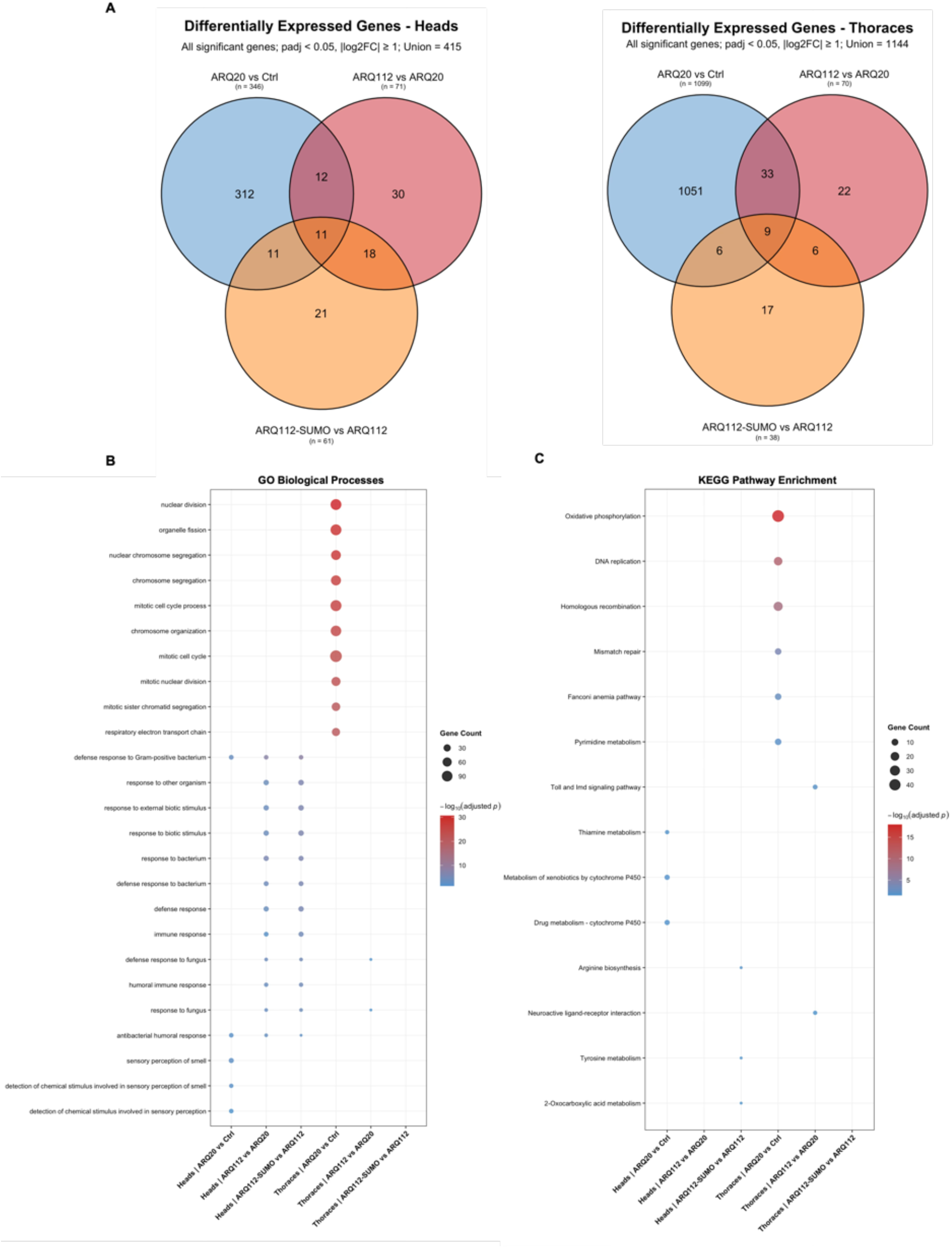
Differential expression and functional enrichment in heads and thoraces of SBMA flies. A) Venn Diagrams showing the number and overlap of differentially expressed genes among comparisons in heads and thoraces. Supplemental tables 1 and 2 provide detailed lists of all DEGs across comparisons. B) GO biological processes enrichment of DEGs. Dot size represents gene count and color indicates enrichment significance (–log10 adjusted p-value). C) KEGG pathway enrichment of DEGs. Dot size represents gene count and color indicates enrichment significance (–log10 adjusted p-value).

**Figure 7:**
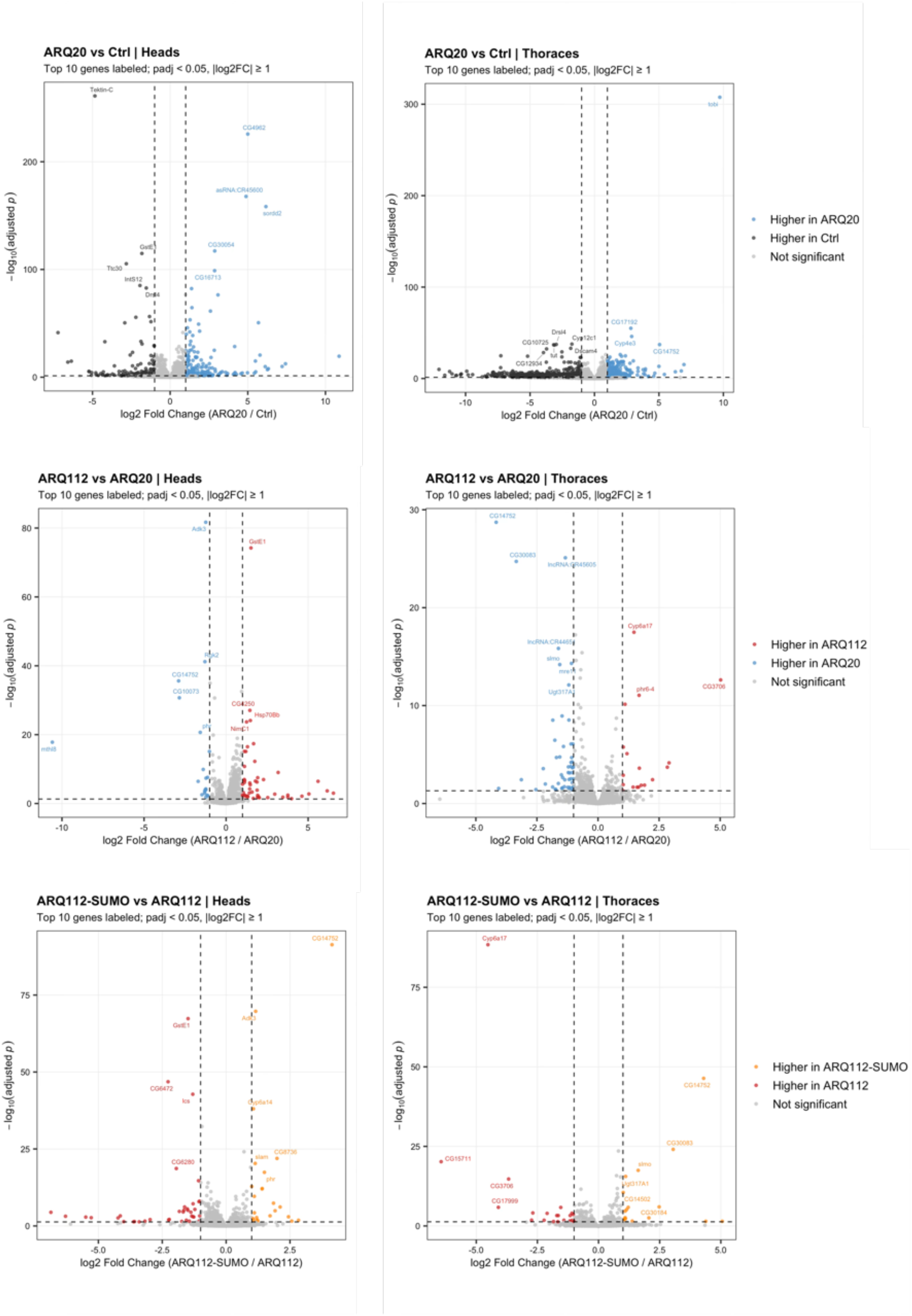
Volcano plots of transcriptional changes across genotypes and tissues. Volcano plots from differential gene expression in heads (left) and thoraces (right) highlighting differences across genotype comparisons. Each point represents an individual gene, with colored points indicating significantly differentially expressed genes (adjusted p < 0.05 and |log2FC| ≥ 1). Color distinguishes genes expressed at higher levels in either genotype within each comparison. The top 10 genes based on significance are labeled in each plot.

Since we were particularly interested in the transcriptional changes associated with disruption of AR SUMOylation in the context of polyQ-expanded AR, we more closely examined the significant DEGs from the ARQ112-SUMO versus ARQ112 volcano plots individually (Figure 8). Consistent with the limited overlap observed between tissues in figure 6, most ARQ112-SUMO-associated DEGs were specific to either heads or thoraces. In heads, increased DEGs included several cytochrome P450 genes (Cyp6a14, Cyp6a20, and Cyp6a23), as well as genes associated with immune-related melanization (PPO1 and PPO2), DNA repair (phr), and sensory processes (Ir51a, Obp56c, and Obp99b). Decreased DEGs included several associated with immune and antimicrobial responses, including DptB, Dro, Drs, AttA, and Mtk, as well as genes involved in transposon regulation and genome stability (piwi and rhi) and detoxification (GstE1). In thoraces, increased DEGs included those associated with fatty-acid metabolism (hll), mitochondrial function (slmo), glycosylation (Ugt317A1), and cell-cycle regulation (CycB). Decreased DEGs included several cytochrome P450 genes (Cyp6a16, Cyp6t3, and Cyp6a17), as well as genes associated with immune function (NimC1), transcriptional regulation (Fer3), and proteolysis (Ser6).

**Figure 8:**
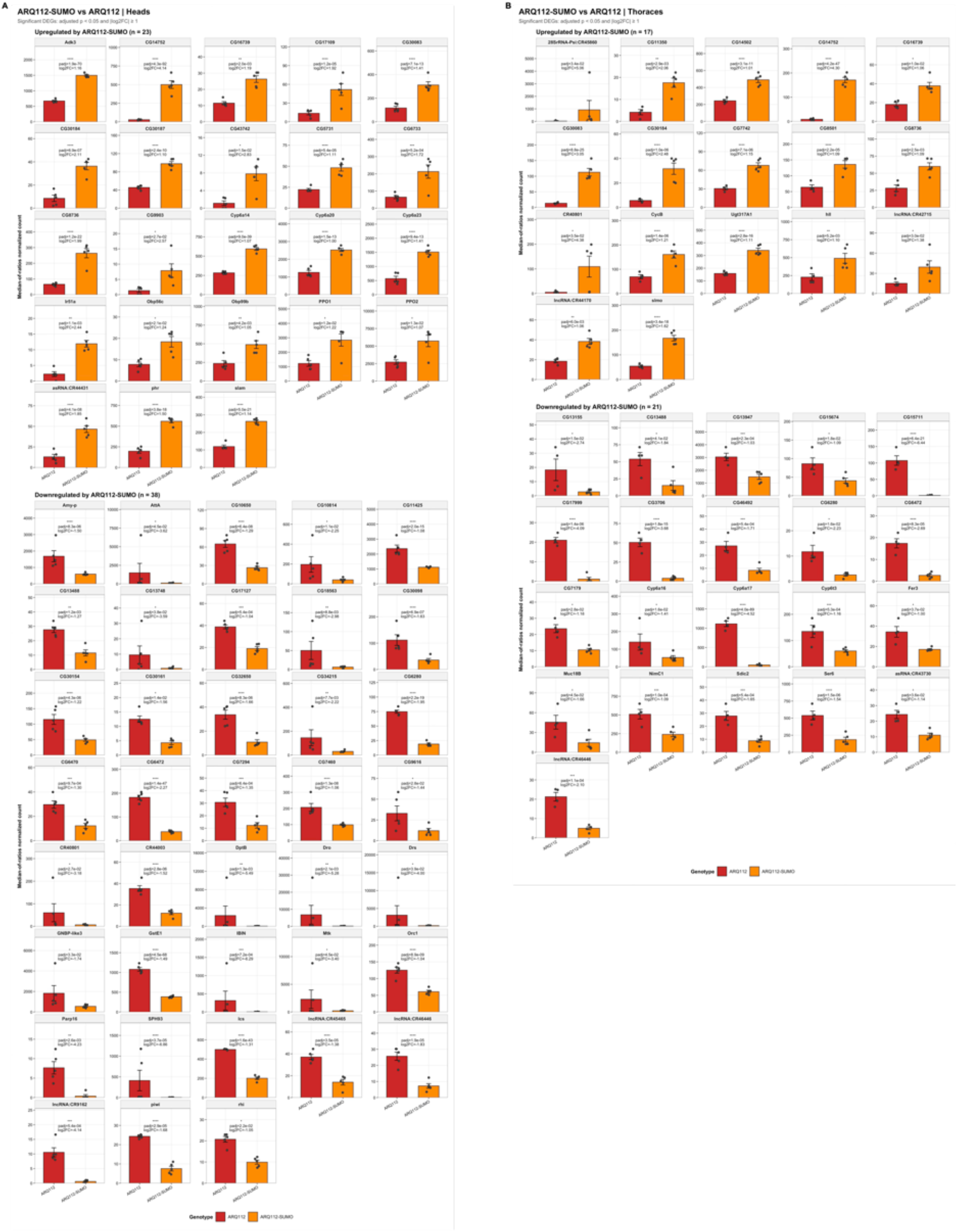
Summary of individual differentially expressed genes following ARQ112-SUMO expression in heads and thoraces. Bar graphs showing normalized counts for significantly differentially expressed genes between ARQ112 and ARQ112-SUMO in heads (A) and thoraces (B). Genes are listed in alphabetical order and separated by upregulation (top) or downregulation (bottom) in ARQ112-SUMO relative to ARQ112.

**Figure 9:**
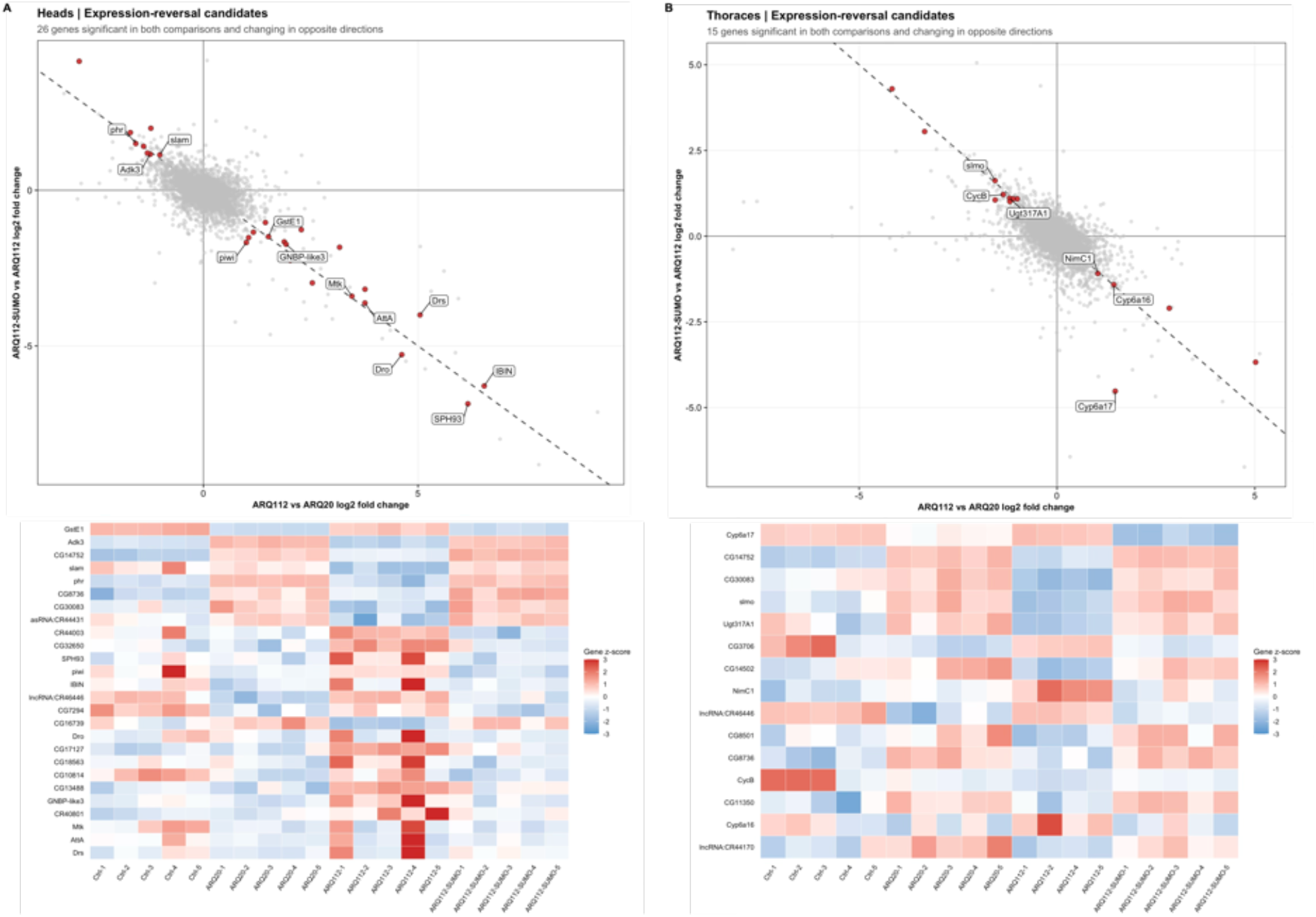
Reversal of polyQ-associated transcriptional changes with ARQ112-SUMO expression. Scatterplots comparing log2FC values from ARQ112 versus ARQ20 and ARQ112-SUMO versus ARQ112 in heads (A; top) and thoraces (B; top). Each point represents an individual gene, with red points indicating genes significantly differentially expressed in both comparisons and changed in opposing directions. Select genes with known characterization are labeled for ease in discussion. Heatmaps (A and B, bottom) show row-scaled log2 normalized expression patterns of all significant genes identified in top panels.

Despite most of the changes being tissue-specific, 10 genes were differentially expressed in both tissues. Of these, nine changed in the same direction. CG14752, CG30083, CG30184, CG8736, and CG16739 were increased in both heads and thoraces, while CG6472, lncRNA:CR46446, CG6280, and CG13488 were decreased in both. The only shared DEG that changed in opposite directions was the pseudogene CR40801, which was decreased in heads but increased in thoraces. Notably, the shared DEGs were largely composed of poorly characterized genes and non-coding transcripts.

### ARQ112-SUMO expression largely reverses polyQ-associated transcriptional changes

Given that a subset of genes altered by ARQ112-SUMO expression overlapped with those altered by ARQ112, we next asked whether disruption of AR SUMOylation further enhanced or opposed the transcriptional changes associated with polyQ expansion. In heads, 26 of the 29 genes shared between the ARQ112 versus ARQ20 and ARQ112-SUMO versus ARQ112 comparisons changed in opposing directions (Figure 9A). Similarly, in thoraces, all 15 shared DEGs changed in opposing directions (Figure 9B). In total, 41 of the 44 overlapping DEGs across both tissues changed in the opposite direction, indicating that disrupting SUMOylation sites on AR largely shifted polyQ-associated transcriptional changes back toward the expression patterns observed with wild-type AR. In heads, genes showing this reversal pattern included GstE1, Adk3, slam, phr, SPH93 and piwi. In thoraces, reversal was observed for genes including Cyp6a17, Cyp6a16, slmo and Utg317A1. Heatmap analysis further highlighted these opposing expression patterns across individual biological replicates (Figure 9 A and B; bottom) for all 41 genes identified in this analysis.

Together, these findings indicate that ARQ112-SUMO expression induces downstream transcriptional changes in both neurons and in muscle, with specific transcriptional changes differing substantially between tissues. Importantly, among the genes whose expression was altered by both polyQ expansion and disruption of AR SUMOylation, ARQ112-SUMO expression largely reversed the direction of the polyQ-associated transcriptional changes. These results suggest that disrupting AR SUMOylation counteracts some of the transcriptional consequences of polyQ-expanded AR in a tissue-dependent manner, which may contribute to the observed tissue-specific benefits.

## Discussion

SUMOylation is an established regulator of AR function, but its contribution to SBMA pathology is incompletely defined. In the present study, we generated a *Drosophila* model of SBMA expressing a SUMO-site mutant form of polyQ-expanded AR and used this system to investigate how disruption of AR SUMOylation sites influences disease-associated phenotypes and downstream gene expression *in vivo*. We found that despite no detectible changes in AR protein abundance, expression, localization or aggregation, ARQ112-SUMO mutants gained a muscle-specific survival benefit, suggesting that SUMOylation may primarily influence downstream functional consequences of mutant AR. No meaningful changes were observed for motility in any tissue tested and aged SBMA flies generally accumulated higher levels of mutant AR protein. Consistent with previous SUMO-mutant SBMA models^22^, SUMO-site mutation did not significantly alter this pattern. However, trends that did not reach statistical significance were observed in soluble/pellet fractions where aged ARQ112-SUMO expression in heads led to AR protein levels resembling younger flies and ARQ112 expression produced a trend toward a larger pelleted fraction.

Our RNA-seq analysis revealed that the transcriptional consequences of ARQ20, ARQ112, and ARQ112-SUMO differed substantially between neuronal and muscle enriched tissues, with very little overlap. Perhaps unsurprisingly, expression of ARQ20 produced the greatest transcriptional response, particularly in muscle. However, the biological processes associated with these changes differed between tissues. In heads, transcriptional changes were largely associated with immune and defense responses, while thoraces showed enrichment of pathways involved in oxidative phosphorylation, DNA replication and repair, and cell-cycle regulation. Although relatively few genes were differentially expressed following ARQ112-SUMO expression in either tissue, and no significant biological process or pathway enrichment was identified in thoraces, several of the DEGs identified have known roles in cellular stress responses, metabolism, and genome maintenance. These findings suggest that AR SUMOylation may influence multiple downstream processes, with effects that are tissue dependent.

Although we were unable to directly demonstrate SUMO protein conjugation to human AR by endogenous *Drosophila* SUMO or confirm loss of this modification in the mutant lines, several observations support the functional relevance of the introduced mutations. First, sequencing verified that the only alteration to the AR transgene between polyQ AR and polyQ AR-SUMO conditions was the substitution of the two established SUMO acceptor lysine residues. Second, expression of ARQ112-SUMO in muscle led to a reproducible extension of lifespan despite no detectable changes in AR transcript or protein levels, indicating that the mutations altered AR function in a biologically meaningful manner. Still, the inability to directly assess AR SUMOylation in our model represents an important limitation of the current study. Detection of SUMOylated proteins in *Drosophila* remains technically challenging because of the limited availability of reagents for endogenous Smt3. Previous studies have frequently addressed this limitation using tagged or modified SUMO constructs^34,42,43^. However, we intentionally avoided introducing exogenous SUMO transgenes into our model. Because SUMOylation regulates a broad range of cellular processes, altering the endogenous SUMO pool could introduce secondary effects that complicate interpretation of AR-specific phenotypes. Instead, we chose to preserve endogenous SUMO and directly modify the SUMO acceptor sites within AR itself. While this approach limits our ability to biochemically verify endogenous SUMO conjugation to our exogenous AR protein, it allows phenotypic changes to be interpreted primarily in the context of AR rather than broader perturbations of the SUMOylation machinery.

One of the most notable phenotypic findings from this study is the tissue specificity of the observed longevity benefit. SUMO-site mutation of AR improved longevity when expressed in muscle but not when expressed in neurons. Increasing evidence suggests that skeletal muscle is not simply a downstream target of motor neuron degeneration in SBMA but is itself a primary site of toxicity that actively contributes to disease progression^44–46^. Our results further support the importance of muscle as a critical mediator of SBMA pathogenesis and suggest that SUMO-dependent regulation of AR is particularly relevant in this tissue.

The mechanism underlying this muscle-specific benefit remains unclear. Given the established role of SUMOylation as a transcriptional repressor of AR activity, one possibility is that preventing SUMOylation partially restores AR-mediated transcriptional programs that become dysregulated in disease. Previous work demonstrated that mutation of AR SUMO acceptor sites enhances transcriptional activity of polyQ-expanded AR and normalizes expression of subsets of androgen-responsive genes^22^. Consistent with this model, we did not observe changes in AR protein levels or aggregation that could account for improved longevity, suggesting that altered downstream signaling rather than altered protein turnover drives the observed protective effect.

One of the most interesting findings from our sequencing results was that ARQ112-SUMO expression opposed many of the transcriptional changes associated with polyQ expansion. Of the 44 genes differentially expressed in both the ARQ112 versus ARQ20 and ARQ112-SUMO versus ARQ112 comparisons, 41 changed in opposing directions, including 26 of 29 genes in heads and all 15 genes in thoraces. However, the interpretation of these changes is complicated by the fact that AR is exogenously expressed in our fly model. Given that AR functions as a transcription factor and flies do not normally express AR, transcriptional changes following its introduction are expected. Indeed, for many of the genes showing a reversal pattern, ARQ20 initially altered gene expression relative to the no-AR control, while ARQ112 shifted expression back toward control patterns. This suggests that polyQ expansion disrupts some of the transcriptional functions associated with wild-type AR, consistent with a potential loss-of-function component. However, despite shifting the expression of these genes closer to the no-AR control, ARQ112 is associated with substantially greater toxicity than ARQ20 in our model^36^. At a broader level, this suggests that the transcriptional changes captured by our analysis do not fully account for the toxicity associated with polyQ-expanded AR. It is possible that more subtle changes in the expression of individual genes, which do not reach statistical significance, collectively contribute to toxicity through broader disruptions in cellular pathways. Additionally, our analysis does not capture changes at the post-translational level, which may contribute substantially to the toxic effects of ARQ112.

In heads, the transcriptional response was strongly associated with immune signaling, defense responses, and detoxification. ARQ112-SUMO expression increased several cytochrome P450 genes, including Cyp6a14, Cyp6a20, and Cyp6a23, as well as the melanization-associated genes PPO1 and PPO2. In contrast, several antimicrobial peptide genes, including DptB, Dro, Drs, AttA, and Mtk, showed decreased expression. These opposing changes suggest that disruption of AR SUMOylation does not simply increase or suppress immune-related transcription but alters specific components of the response. This is particularly interesting given the growing evidence that dysregulated immune responses contribute to neurodegenerative diseases, including polyQ disorders^46,47^. However, it remains unclear whether the changes observed here represent protective responses, altered immune regulation, or secondary consequences of AR expression.

In thoraces, a significant enrichment of oxidative phosphorylation was observed following ARQ20 expression relative to control. Mitochondrial dysfunction and impaired energy metabolism have been implicated in several neurodegenerative diseases, including SBMA^45,46,48,49^. Oxidative phosphorylation was not significantly enriched in the ARQ112 versus ARQ20 or the ARQ112-SUMO versus ARQ112 comparisons using our initial DEG-based enrichment analyses. However, further investigation using targeted gene sets revealed a strong negative enrichment of oxidative phosphorylation and aerobic respiration in ARQ112 versus ARQ20 and a strong opposing pattern of positive enrichment in ARQ112-SUMO, specifically in thoraces but not in heads (Supplemental figure 10). These findings suggest that AR SUMOylation influences the transcriptional regulation of pathways involved in mitochondrial energy production and raise the more narrow possibility that altered energy metabolism may contribute to the protective effects on lifespan observed in muscle.

An important consideration when studying human disease in preclinical models is whether the findings are relevant to disease mechanisms in mammals and in humans. In a mouse model of SBMA, replacement of the conserved lysine resuidues that serve as SUMO acceptor sites enhanced the transcriptional activity of polyQ-AR and rescued exercise endurance and type I muscle fiber atrophy while prolonging survival; these benefits occurred without overt alteration in polyQ-AR expression or aggregation^22^. Our observations in *Drosophila* recapitulate this pattern closely: ARQ112-SUMO flies gained a muscle-specific survival benefit in the absence of detectable changes in AR abundance, subcellular localization or aggregation, and the accompanying transcriptional changes largely opposed those associated with polyQ expansion.

Although *Drosophila* muscle does not have the same fiber-type organization of mammalian muscle, the indirect flight musculature is highly oxidative^50^. This point is pertinent given that the rescue observed in mouse models was evident in oxidative type I fibers. That SUMOylation acts on polyQ-AR through routes separable from aggregation is further supported by evidence that enhancing SUMOylation reduces polyQ-AR aggregate formation without altering transcriptional activity^25^ and by earlier work in flies where perturbing the SUMO-1 conjugation pathway modulated polyQ-AR-induced degeneration^24^. Together with evidence that skeletal muscle is a major site of toxicity in SBMA^44,46,51^, these parallels suggest that the muscle-restricted benefit we observed reflects a conserved, SUMO-dependent modulation of mutant AR function rather than a change in its disposition.

In summary, our findings suggest that AR SUMOylation influences SBMA pathology in a tissue-dependent manner, potentially through changes in downstream gene expression rather than discernible AR protein accumulation. Although the effects of these transcriptional changes at the protein level are unknown, our results add to growing evidence that SUMOylation plays a complex and context-dependent role in SBMA and further highlight the importance of peripheral tissues in disease. Future studies will be important for identifying the mechanisms through which SUMOylation influences mutant AR toxicity and determining whether these pathways may provide therapeutic benefit in SBMA.

## Methods

### Fly Stocks and Maintenance

Flies were maintained in temperature and humidity-controlled incubators, at 25°C on a 12-hour light-dark cycle for the duration of the experiments. Virgin females and males collected for crosses and used in experiments were sorted under light CO_2_ anesthesia. To ensure age matching, all flies used in experiments were collected within the same 72-hour window. Flies were flipped to fresh food vials every 2 days to maintain healthy conditions. Unless otherwise specified, all experiments were conducted using female flies. The following stocks were used: *elav*-Gal4 (X chromosome; Bloomington Drosophila Stock #458), *Mef2*-GAL4 (chromosome 3; Bloomington Drosophila Stock #27390), and lab generated UAS constructs UAS-ARQ112, UAS-ARQ112-SUMO (chromosome 3).

### Antibodies

#### Primary antibodies

rabbit anti-AR (Cell Signaling Technology, Catalog #: 5153S), rabbit anti-Smt3/SUMO (1:1000, ThermoFisher Scientific, Catalog #: PA-571762), mouse anti-Lamin (1:50, Developmental Studies Hybridoma Bank, Catalog #: AB-528338), mouse anti-Tubulin (1:1000, Sigma Aldrich, Catalog #: T5168). *Secondary antibodies:* goat anti-mouse horseradish peroxidase (1:2500, Jackson ImmunoResearch Laboratories, Catalog #: 115-035-146), goat anti-rabbit horseradish peroxidase (1:2500, Jackson ImmunoResearch Laboratories, Catalog #: 111-035-144).

### Generation of SUMO Mutants

The full-length human AR sequence used to generate our SBMA fly lines was derived from the NCBI reference sequence NM_000044.6 and codon-optimized for use in *Drosophila*^36^. In this reference sequence, established SUMOylation sites are located at K388 and K521, with a polyQ tract length of 23Q. In the SBMA model, expansion of the polyQ tract to 112Q results in a positional shift of these SUMOylation sites to K478 and K611. These lysine residues were mutated to arginine via A to G nucleotide substitutions to prevent SUMOylation (Supplemental figure 1). pWalium10-moe plasmid constructs were synthesized by Genscript and injected by model systems injections using pHiC31-mediated site-specific integration into VK00033 on chromosome 3. The resulting transgenic lines were subsequently established and maintained in-house.

### Climbing Speed

Climbing ability was assessed in two complementary ways: rapid iterative negative geotaxis (RING) and timed motility. For RING assays, five vials containing 20 flies each (N=100) were flipped into fresh, empty vials and placed in a RING apparatus. Flies were allowed to equilibrate to room conditions for approximately 20 minutes prior to assessments. Negative geotaxis was induced simultaneously across all five vials by firmly and rapidly tapping the apparatus on the countertop. A digital image was taken 2 seconds after the initiation of climbing response. Images were taken three times per week at a consistent time of day to track longitudinal changes in climbing performance. A total of 20 images per week were analyzed to calculate an average climbing index, defined as the mean height reached by flies within their vials after 2 seconds.

For descriptions in further detail, reference Damschroder *et al*., 2018^52^. For timed motility assays, five vials containing 10 flies per group were placed in the RING apparatus and allowed to equilibrate for 20 minutes prior to stimulating negative geotaxis. Once stimulus was applied, 30-second videos were recorded to capture continuous climbing data. Assessments took place once per week over a 5-week period. Videos were analyzed manually by tracking the number of flies that reached the top within 5, 15 and 30 seconds after stimulation. The percent of flies that reached the top was calculated for each vial and averaged across all five vials to obtain the mean percentage of flies reaching the top of the vial within each time window.

### Longevity

Survival curves were generated using cohorts of 200 flies per condition. All cohorts were age matched, and death events were recorded five days per week throughout the lifespan at the time of vial flipping. Flies that escaped the vials during handling or died from non-aging related causes (e.g., accidental injury during transfer) were excluded from the analysis.

### Quantitative PCR

Relative AR expression in *Mef2* ARQ112 and ARQ112-SUMO conditions was measured using qrt-PCR analysis. RNA was isolated by homogenizing 20 whole flies per genotype in ice cold TRIZOL (Invitrogen) under RNAse free conditions. The resulting RNA was converted to cDNA using a High-Capacity cDNA Reverse Transcription Kit (Applied biosystems, Catalog #: 4368814) following the manufacturer’s protocol. A total of three biological reps per condition and four technical reps were loaded onto a 96-well plate. Each well contained *Power*SYBER Green PCR Master Mix (10uL; Applied biosystems, Catalog #: 4367659), cDNA (2uL at 5ng/uL concentration), forward and reverse primers (2uL at 5uM concentration) and nuclease-free water (6uL). Plates were run on a QuantStudio 5 Real-time PCR system (Thermo Fisher Scientific) and resulting mRNA data were quantified using cycle threshold methods and normalized to endogenous rp49 expression. Primers used were as follows:

rp49-F: 5′-AGATCGTGAAGAAGCGCACCAAG-3′

rp49-R: 5′-CACCAGGAACTTCTTGAATCCGG-3′

AR-F: 5′-GATATGAGGCTGGAGACCGC-3′

AR-R: 5′-ACAGGTACTTCTGCTTGCCC-3′

### Co-Immunoprecipitations

30 whole flies were homogenized in 1mL of NETN lysis buffer (50mM Tris, pH 7.5, 150 mM NaCl, 0.5% Nonidet P-40) supplemented with protease inhibitors (Millipore Sigma, Catalog #: S-8820). Samples were incubated at 4°C on rotation for 30 minutes and centrifuged at 5000 xg for 5 minutes. Resulting supernatants were combined with HA-bound beads and incubated on rotation at 4°C for four hours. Beads were rinsed five times each with lysis buffer (50mM Tris pH 6.8, 2% SDS, 10% glycerol, 100mM dithiothreitol) and bead-bound complexes were eluted by adding Laemmli buffer (12.5% 1M Tris, 20% Glycerol, 4% SDS, pH7.5) and boiling. Input and post-IP samples were visualized by western blotting.

### Soluble/Pellet Preparations

10 whole flies were homogenized in NETN lysis buffer supplemented with protease inhibitors the morning of lysis. Lysates were sonicated for 15 seconds at 50% amplitude and subsequently centrifuged for 30 minutes at 20,000 xg at 4°C. The resulting supernatant (soluble fraction) and the pellet (insoluble fraction) were separated into new microcentrifuge tubes. The pelleted fractions were resuspended in 50uL Laemmli buffer and sonicated again for 30 seconds. All samples were then supplemented with 6% SDS, boiled for 10 minutes, and centrifuged at room temperature for 7 minutes at 13,300 rpm. Fractions were analyzed by western blotting and quantified as a percentage of the total AR signal [example: soluble/(soluble + pellet) : 100].

### Cytoplasmic/Nuclear Fractionations

Cytoplasmic and nuclear fractions were prepared using the ReadyPrep Protein Extraction Kit (Cytoplasmic/Nuclear; Bio-Rad). Five whole flies per group were homogenized in cytoplasmic extraction buffer (Bio-rad). The isolated nuclear fraction (pellet) was resuspended in Protein Solubilization Buffer following the manufacturer’s instructions. Fractions were subsequently analyzed by western blotting and quantified as a percentage of the total AR signal.

### Western Blotting

For western blot analyses, 5-30 whole flies (where specified for each experiment type) were aged to indicated time points and frozen at -80C prior to use. Preparation of samples analyzed by western blot were performed as follows: For whole fly lysates, five flies were homogenized in boiling lysis buffer, sonicated for 15 seconds at 50% amplitude, boiled for 10 minutes, and centrifuged at 13300rpm for 7 minutes. Supernatants were loaded onto either 7.5% or 4-20% pre-cast gels where indicated in the figure legends and separated by SDS-PAGE. Following electrophoresis, proteins were transferred to a PVDF membrane. Membranes were incubated in primary and secondary antibodies and imaged using a Chemidoc imaging system (Bio-rad).

Total protein staining was performed using Direct Blue 71 as a loading control. PVDF membranes were incubated for 10 minutes with 0.0008 % Direct Blue 71 prepared in 40% ethanol and 10% acetic acid, rinsed with the same solvent followed by water, air dried overnight and imaged on the Chemidoc. Quantifications of western blot and total protein signals were done using ImageLab software.

### Immunohistochemistry

Adult flies were aged and dissected at either 3 or 30 days to reflect early- and late-stage disease, respectively. At each timepoint, tissues were dissected and immunofluorescent staining was completed to visualize AR localization in both brain and muscle tissue, as well as NMJ complexity in the muscle using both neuronal and muscle specific drivers. ***Brain:*** Brain staining protocols were modified from Tito *et al.,* 2016^53^. Adult flies were anesthetized under light CO2 and briefly submerged in 100% ethanol to remove the waxy coating on the cuticle prior to being transferred to a gel-bottomed dissecting dish filled with ice cold 1X Phosphate Buffered Saline (PBS, pH 7.2; Gibco 20012-027). The head casing was gently opened using fine dissecting forceps to expose the underlying brain tissue. Sets of 10 flies with exposed brains were fixed in 4% paraformaldehyde on rotation at room temperature for 1 hour. Samples were then incubated in blocking buffer (1X PBS + 0.3% Triton-X + 2% Bovine Serum Albumin) for 2 hours. Tissues were placed in primary antibody solution overnight at 4°C on rotation. Samples were subsequently incubated in secondary antibodies, followed by staining with DAPI nuclear stain (1:1000, ThermoFisher Scientific, Catalog #: D1306). After staining, tissues were mounted in a 50% glycerol solution and imaged. 20x images were obtained by confocal microscopy and displayed as maximum intensity projections, whereas 100x images were taken as single optical sections. Primary antibodies: rabbit anti-AR (1:200, Cell Signaling Technology, Catalog #: 5153S). Secondary antibodies: goat anti rabbit Alexa Fluor 568 (1:1000, ThermoFisher Scientific, Catalog #: A-11011). ***Muscle:*** Muscle tissue dissections were performed as previously described by Sidisky *et al.,* 2020^54^. Briefly, unconscious flies were submerged in 100% ethanol to remove the wax coating on the cuticle and subsequently transferred to a gel-bottomed dissecting dish containing 1X PBS. Wings, legs and head were removed using dissecting scissors to isolate the thorax. Isolated thoraces were transferred to Eppendorf tubes containing 4% paraformaldehyde and fixed on rotation for 1 hour. Samples were then washed in 1X PBS, flash frozen in liquid nitrogen, and immediately returned to ice in fresh, ice-cold PBS. Individual thoraces were then bisected using a sharp razor blade, washed with 1X PBS + 0.3% Triton-X and incubated in blocking buffer for 2 hours. Primary antibodies were added and samples were incubated overnight on rotation at 4°C. Subsequent secondary antibody staining was completed at room temp for 2 hours, followed by treatment with DAPI nuclear stain and mounting in 50% glycerol before imaging with confocal microscopy. Primary antibodies: rabbit anti-AR (Cell Signaling Technology, Catalog #: 5153S), mouse anti-22c10 (1:50, Developmental Studies Hybridoma Bank, AB-528403). Secondary antibodies: goat anti-rabbit Alexa Fluor 647 (1:500, ThermoFisher Scientific, Catalog #: A-21245), goat anti-mouse Alexa Fluor 488 (1:500, ThermoFisher Scientific, Catalog #: A-11001), Phalloidin Alexa Fluor 594 (1:1000, ThermoFisher Scientific, Catalog #: A-12381).

### RNA sequencing and Differential Expression analyses

For transcriptomic analysis, 70 heads (*elav*) or 20 thoraces (*Mef2*) were isolated from seven-day old female flies and shipped on dry ice to GENEWIZ (South Plainfield, NJ, USA) for RNA extraction, library preparation (N=5 biological replicates per group) and next-generation sequencing. Reads were mapped to the *Drosophila melanogaster* BDGP6 reference genome, and samples were analyzed separately by tissue. For thorax samples, one ARQ112 biological replicate was identified as an outlier and excluded from all subsequent analyses (ARQ112-3; see below for sample exclusion criteria). PCA was therefore performed on 20 head libraries and 19 thorax libraries using the 1,000 most variable genes following normalization and log2 transformation. Genes were filtered to retain those with >20 total raw counts summed across all samples in each pairwise comparison, and library size normalization was performed using DESeq2 median-of-ratios. Differential expression was assessed using DESeq2 Wald tests and p-values were adjusted using the Benjamini–Hochberg method. A gene was considered differentially expressed when the adjusted p-value was <0.05 and |log2FC|≥1. KEGG pathway enrichment analysis was performed using all significant DEGs and *Drosophila melanogaster* pathway annotations from the Kyoto Encyclopedia of Genes and Genomes database. GO enrichment analysis was also performed using all significant DEGs, with GO Biological Process enrichment assessed using the *Drosophila melanogaster* annotation database. For targeted gene set enrichment analysis, genes with >20 total raw counts across samples within each tissue were retained and ranked using the DESeq2 Wald statistic, rather than restricting the analysis to significant DEGs. Enrichment was assessed using GO Biological Process terms, and oxidative phosphorylation and aerobic respiration were selected for visualization. Normalized enrichment scores were used to determine the direction and magnitude of enrichment, with adjusted p-values <0.05 considered significant. Gene sets containing 10–500 genes were included in all enrichment analyses. All resulting graphs were generated using R version 4.6.0^55^.

### Outlier assessment and sample exclusion

Quality control analysis of tissue-separated samples identified a potential outlier in the muscle dataset. To assess this sample prior to downstream analyses, all muscle libraries were evaluated in a formal multi-metric audit. Muscle-library genes were filtered at a count of at least 10 in at least five samples, followed by median-of-ratios normalization and log2 transformation. For each sample, within-genotype Pearson correlations were summarized, including the mean correlation with its two nearest genotype-matched replicates. Multivariate sample distance was assessed across PC1–PC5 using the Euclidean distance from the within-genotype component-wise median after scaling each component by its median absolute deviation (MAD; constant = 1). Sample influence was additionally evaluated using Cook’s distance from a group-means linear model fitted to the log2-normalized expression matrix, with the proportion of filtered genes exceeding a Cook’s distance of 4/N recorded for each sample, where N represents the number of muscle libraries. Muscle ARQ112 sample 3 was the only isolated sample supported by all three measures to be an outlier. Its mean correlation with the two nearest ARQ112 replicates was 0.939, its robust PC1–PC5 distance was 20.3, and its Cook’s distance exceeded 4/N for 59.4% of filtered genes. Based on the concordance of these complementary diagnostics, Muscle ARQ112 sample 3 was excluded before application of the project-specific gene-count filter (>20 counts in at least five samples to match GENEWIZ processing of neuronal samples), normalization, and all other downstream differential-expression analyses.

### Statistics

Statistics were completed and data were visualized using GraphPad Prism version 11. Detailed statistical analyses are included within the figure legends where applicable.

## Acknowledgments

This study was partially supported by NIH grant R01NS086778 (SVT).

## Data Availability

All relevant data are included in the figures, tables and supplemental material. RNA-sequencing dataset will be deposited in NCBI’s Gene Expression Omnibus.

## Author Contributions

KR: Conceptualization, methodology, software, validation, formal analysis, investigation, data curation, writing – original draft, writing – review and editing, visualization.

MVP: Software, validation, formal analysis, data curation, writing – review and editing, visualization.

TK: Validation, investigation, writing – review and editing, visualization. FHH: Validation, investigation, writing – review and editing, visualization.

SVT: Conceptualization, methodology, software, validation, formal analysis, investigation, resources, data curation, supervision, funding acquisition, writing – original draft, writing – review and editing, visualization.

## Declarations

The authors do not have any conflicts of interest to declare.

## Supplemental Materials

**Supplemental figure 1:**
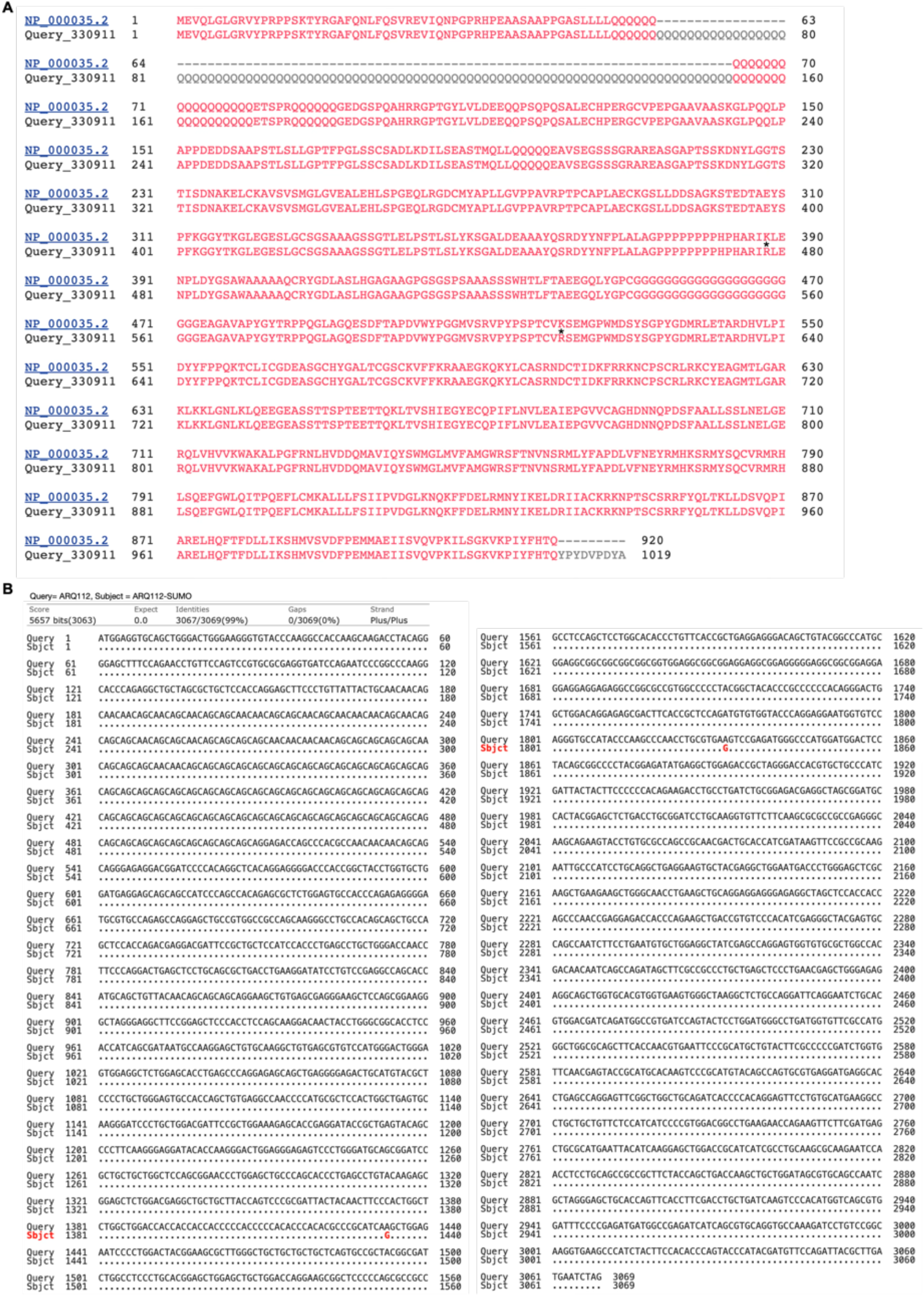
A) Amino Acid sequence alignment of ARQ112 and ARQ112-SUMO highlighting SUMO-site mutations. B) Nucleotide sequence alignment of ARQ112 and ARQ112-SUMO, with the nucleotide substitutions corresponding to the SUMO-site mutations highlighted in red.

**Supplemental figure 2:**
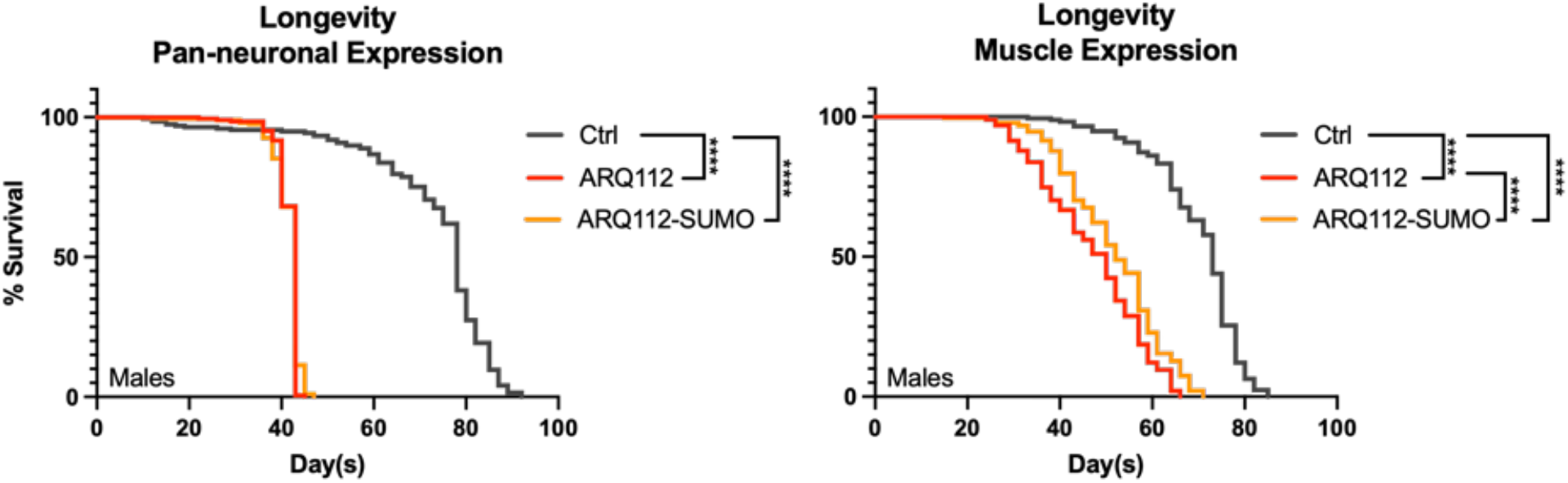
Effect of ARQ112-SUMO expression on longevity in male flies. Longevity outcomes with expression of noted transgenes in neurons (left) and muscle (right). For both panels, N=100 flies per group. Statistics: Log-rank tests, ****: p<0.0001.

**Supplemental figure 3:**
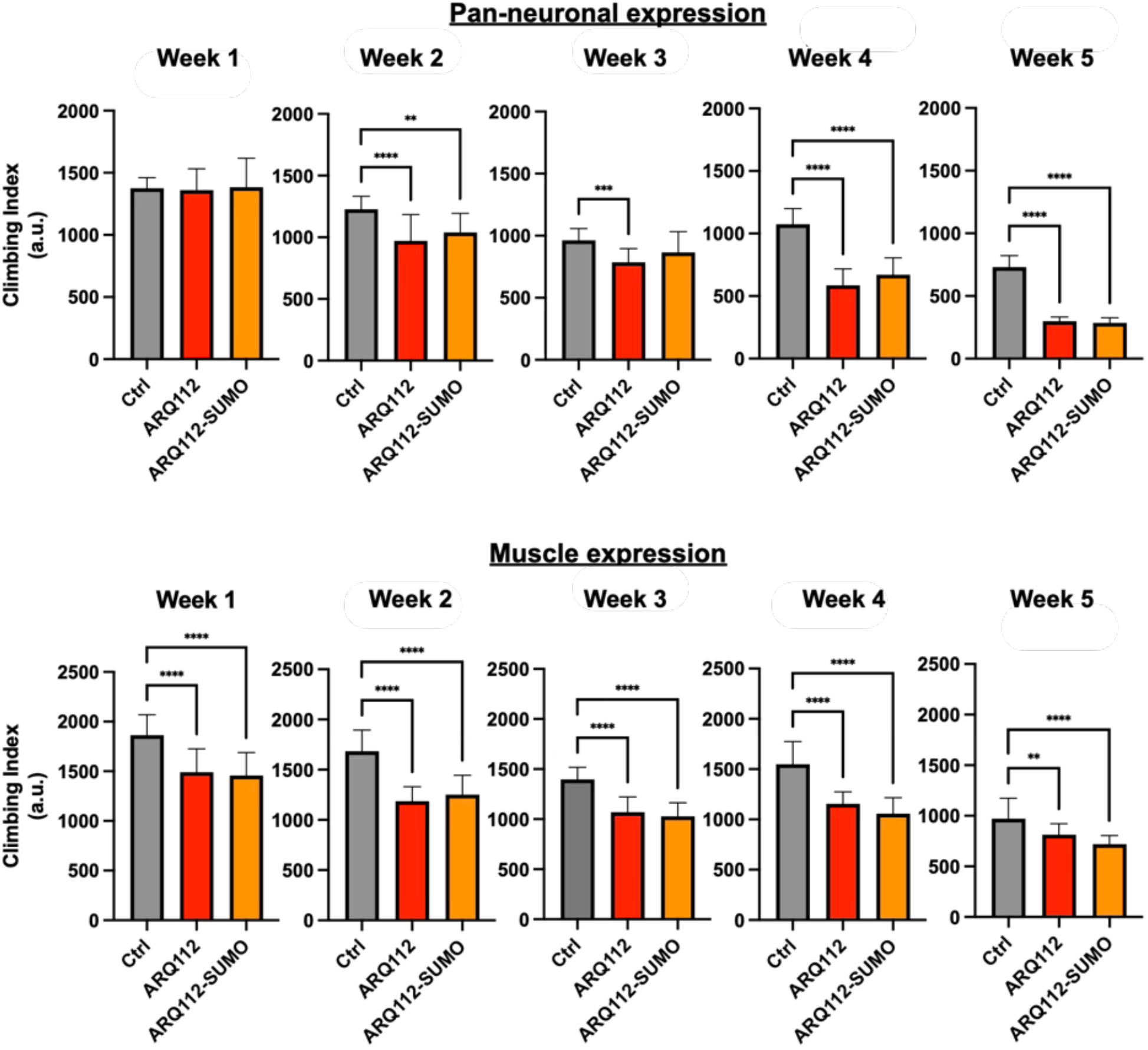
Effect of ARQ112-SUMO expression on motility by week. Climbing ability of flies expressing the noted transgenes in neurons (top) or muscle (bottom) measured weekly. N=100 flies per group. Statistics: One-way ANOVA + Tukey post-hoc, mean ± SD. **: p<0.01, ***: p<0.001, ****: p<0.0001.

**Supplemental figure 4:**
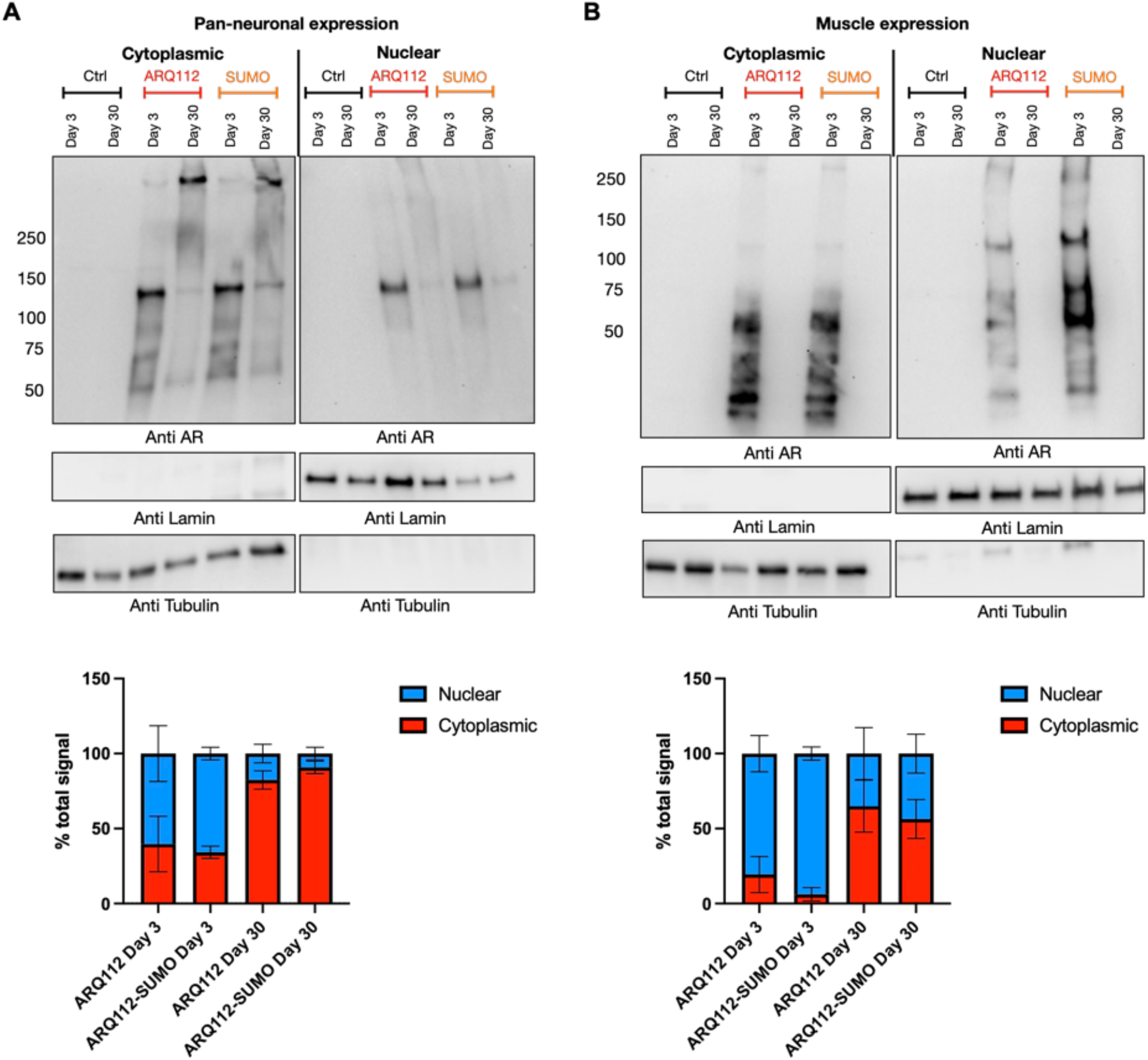
Subcellular localization of ARQ112 and ARQ112-SUMO in neurons and muscle. A) Western blots of cytoplasmic and nuclear fractions from flies expressing ARQ112 or ARQ112-SUMO in neurons at day 3 and day 30. N=3 independent lysates per condition. Blots were run on 4-20% gels and quantified as a percentage of the total AR signal. Lamin and tubulin were used as nuclear and cytoplasmic fraction markers, with AR signal normalized to total protein within each fraction. Statistics: not significant by unpaired Welch’s t-test, mean ± SD. B) Western blots of cytoplasmic and nuclear fractions from flies expressing ARQ112 or ARQ112-SUMO in muscle at day 3 and day 30. N=3 independent lysates per condition. Blots were run on 4-20% gels and quantified as a percentage of the total AR signal. Lamin and tubulin were used as nuclear and cytoplasmic fraction markers, with AR signal normalized to total protein within each fraction. Statistics: not significant by unpaired Welch’s t-test, mean ± SD.

**Supplemental figure 5:**
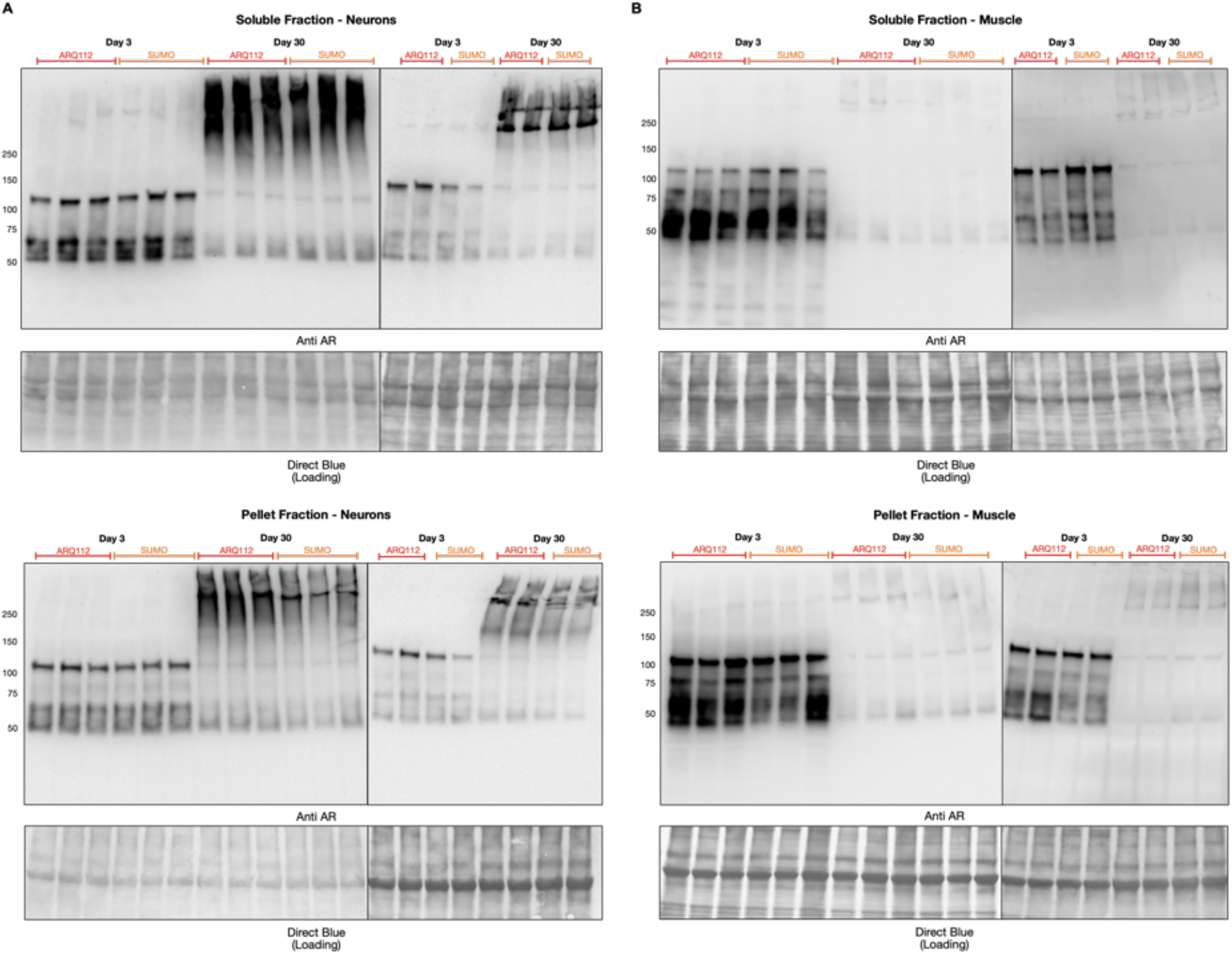
Soluble and pellet fractionation of ARQ112 and ARQ112-SUMO in neurons and muscle. Western blots showing AR protein in soluble (top) and pellet (bottom) fractions from flies expressing ARQ112 or ARQ112-SUMO in neurons (A) or muscle (B) at days 3 and 30. All five biological replicates (N = 5) used for the quantification in figures 4D and 5C are shown.

**Supplemental figure 6:**
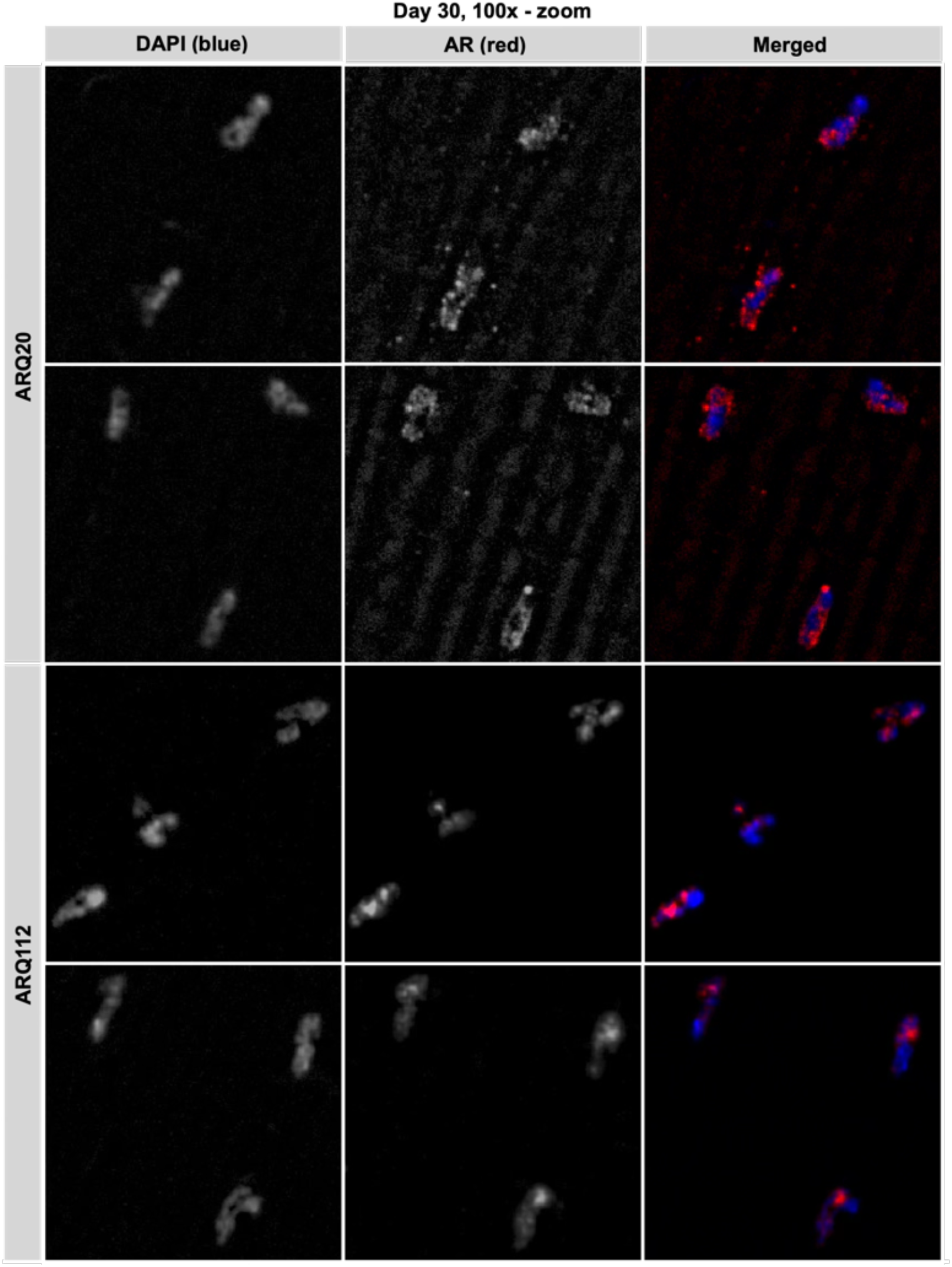
AR protein localization of ARQ20 and ARQ112 expressed in muscle. Representative immunofluorescence images showing AR protein localization with ARQ20 or ARQ112 expression in muscle at day 30. 100x magnification, red: anti AR, blue: DAPI. Single channel images are shown in greyscale for ease of visualization.

**Supplemental figure 7:**
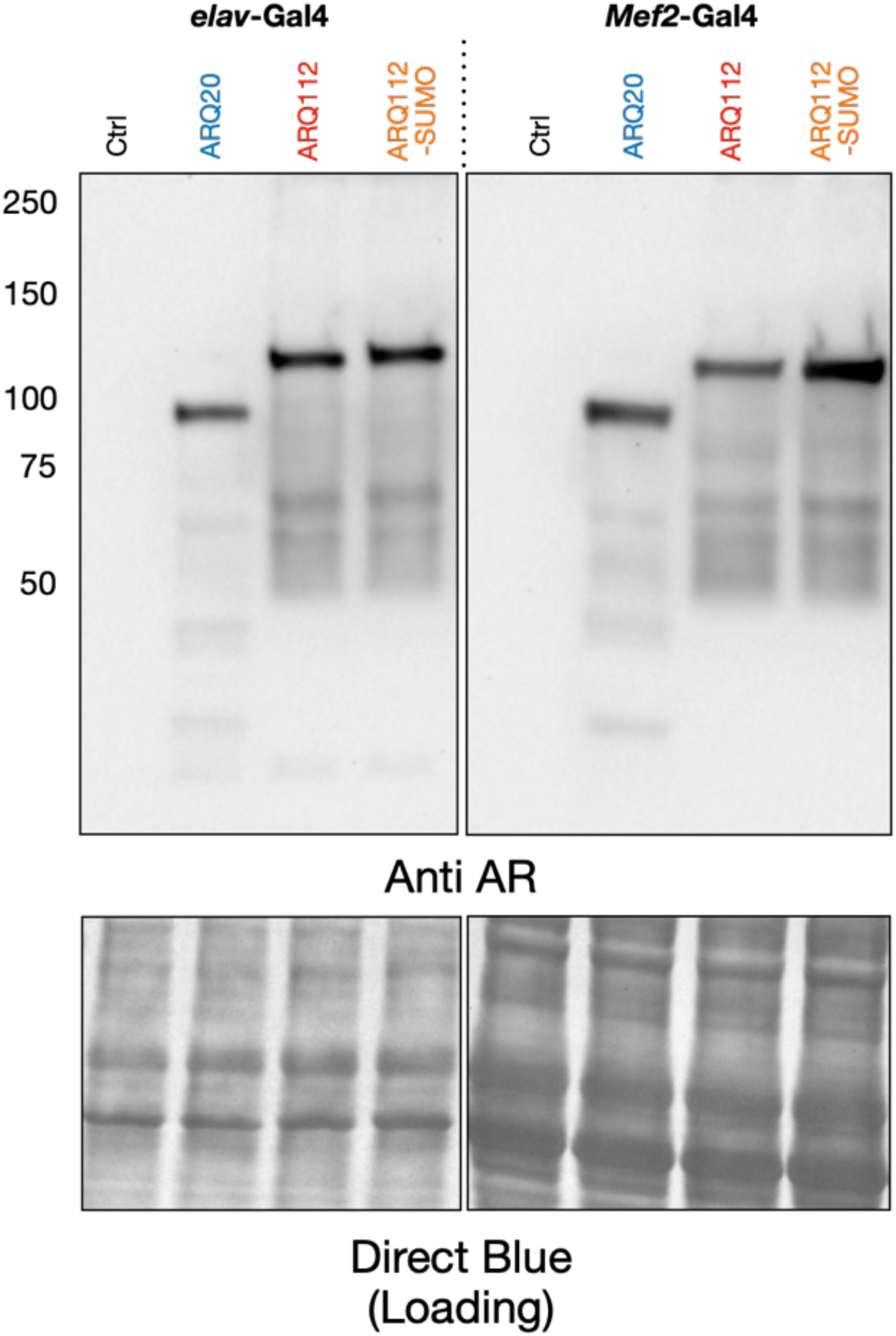
Verification of AR expression in RNA-sequencing samples. Western blot confirming expression of correct AR transgenes in samples sent for RNA-sequencing.

**Supplemental figure 8:**
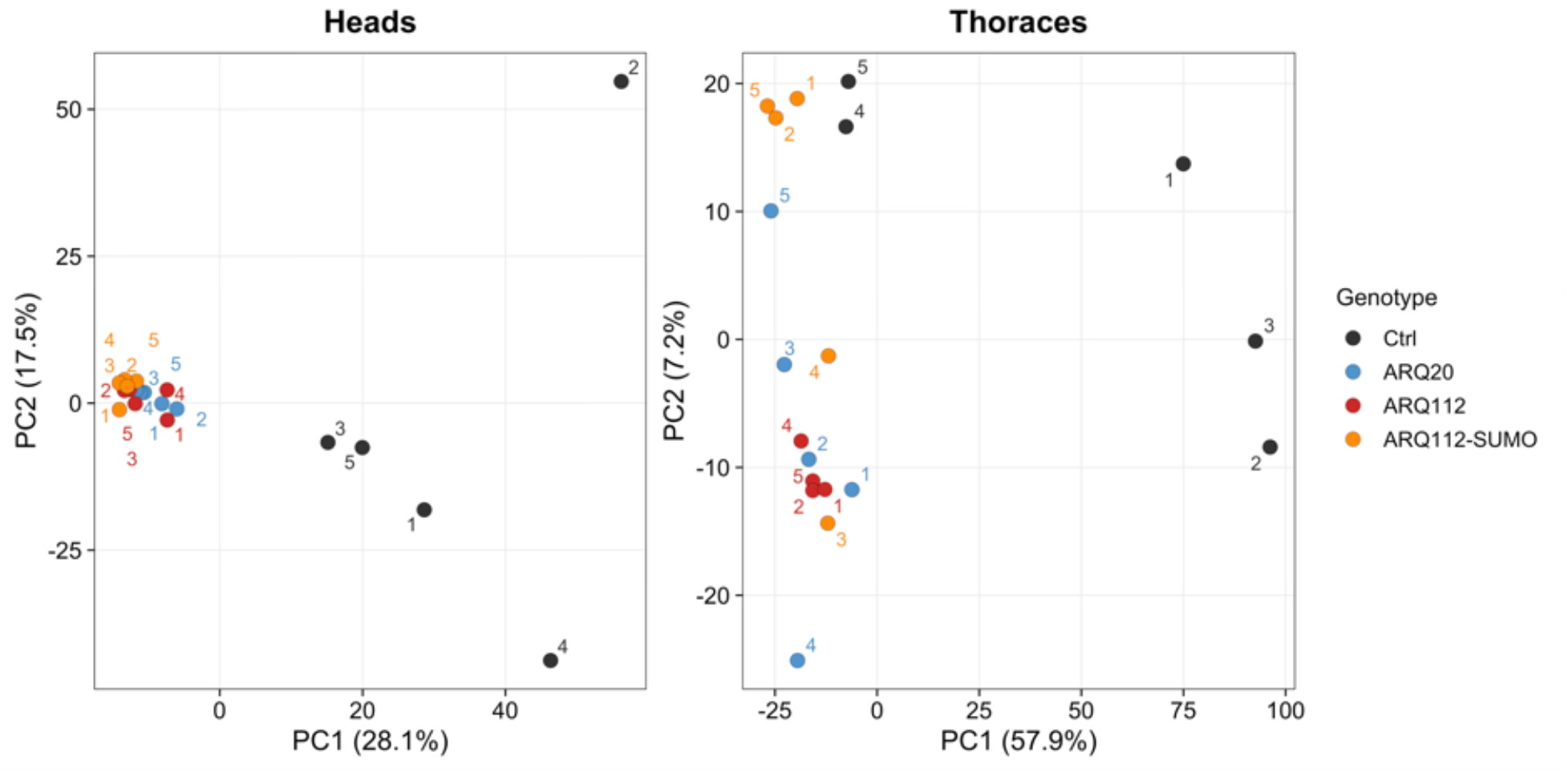
Principal component analysis of RNA-seq samples in heads and thoraces. Principal component analysis of RNA-seq samples from heads (left) and thoraces (right) following outlier exclusion. Each point represents an individual biological replicate, colored by genotype and labeled by replicate number.

**Supplemental figure 9:**
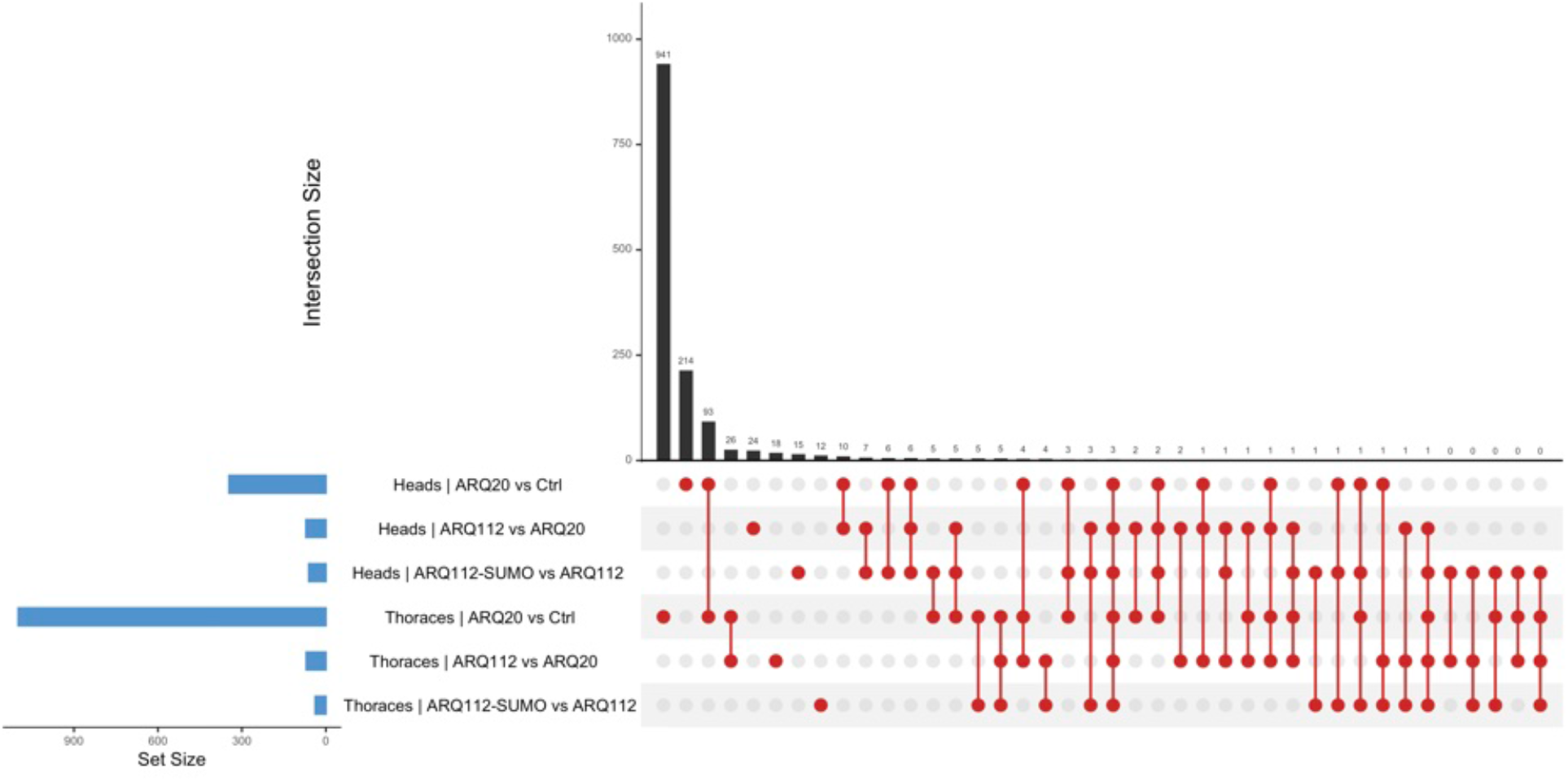
Overlap of differentially expressed genes across RNA-seqcomparisons. UpSet plot showing the number and overlap of differentially expressed genes across comparisons and in both tissues. Blue bars represent the total number of DEGs in each comparison. Black bars indicate the number of genes within each intersection.

**Supplemental figure 10:**
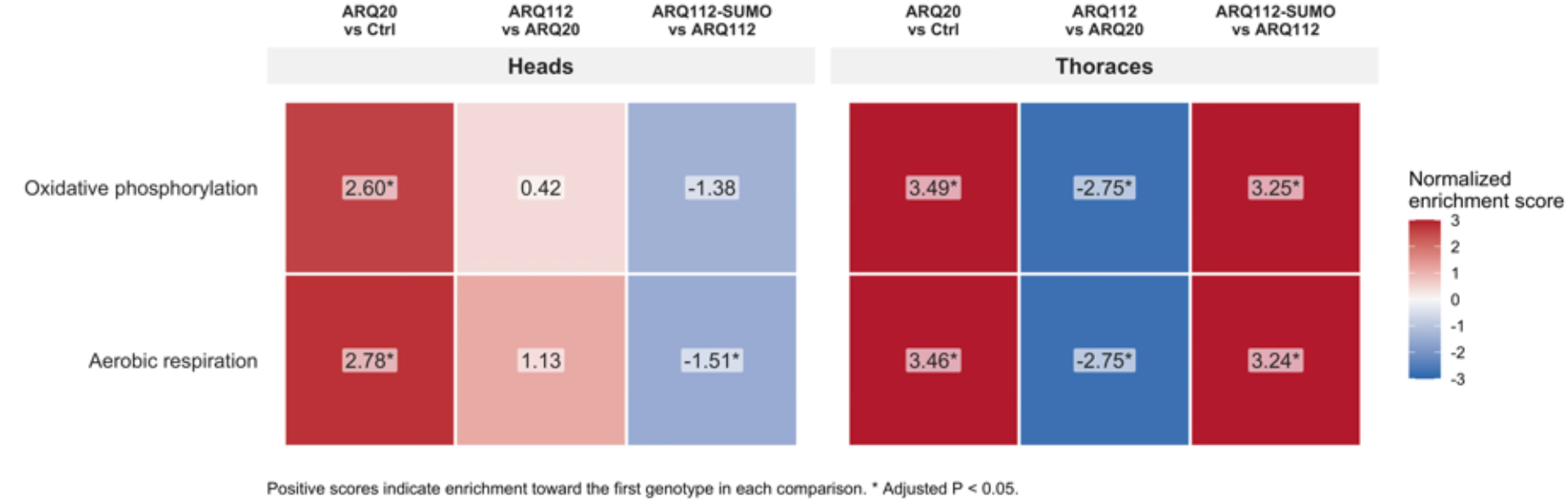
Targeted gene set enrichment analysis for oxidative phosphorylation and aerobic respiration in heads and thoraces. Heatmap showing normalized enrichment scores for oxidative phosphorylation and aerobic respiration gene sets across genotype comparisons in heads (left) and thoraces (right). Positive NES values indicate enrichment toward the first genotype in each comparison, while negative values indicate enrichment toward the second genotype. Colors represent the direction and magnitude of enrichment. Asterisks indicate statistically significant enrichment (adjusted p < 0.05).

Supplemental Table 1: Venn membership of all significant differentially expressed genes in heads

Supplemental Table 2: Venn membership of all significant differentially expressed genes in thoraces

Supplemental Video: Negative Geotaxis induced climbing ability

Video of day 35 negative-geotaxis induced climbing ability with expression of driver-only control (blue or green; w1118), ARQ112 (orange; AR3) or ARQ112-SUMO (yellow; SUMO) in neurons (left) and muscle (right).

